# Magnetically Controlled Cyclic Microscale Deformation of *In Vitro* Cancer Invasion Models

**DOI:** 10.1101/2023.03.30.534990

**Authors:** D.O. Asgeirsson, A. Mehta, N. Hesse, A. Scheeder, R. Ward, F. Li, M. G. Christiansen, A. J. De Micheli, E. S. Ildic, N. Aceto, S. Schuerle

## Abstract

Mechanical cues play an important role in the metastatic cascade of cancer. Three-dimensional (3D) tissue matrices with tunable stiffness have been extensively used as model systems of the tumor microenvironment for physiologically relevant studies. Tumor-associated cells actively deform these matrices, providing mechanical cues to other cancer cells residing in the tissue. Mimicking such dynamic deformation in the surrounding tumor matrix may help clarify the effect of local strain on cancer cell invasion. Remotely controlled microscale magnetic actuation of such 3D *in vitro* systems is a promising approach, offering a non-invasive means for in situ interrogation. Here, we investigate the influence of cyclic deformation on tumor spheroids embedded in matrices, continuously exerted for days by cell-sized anisotropic magnetic probes, referred to as µRods. Particle velocimetry analysis revealed the spatial extent of matrix deformation produced in response to a magnetic field, which was found to be on the order of 200 µm, resembling strain fields reported to originate from contracting cells. Intracellular calcium influx was observed in response to cyclic actuation, as well as an influence on cancer cell invasion from 3D spheroids, as compared to unactuated controls. Localized actuation at one side of a tumor spheroid tended to result in anisotropic invasion toward the µRods causing the deformation. In summary, our approach offers a strategy to test and control the influence of non-invasive micromechanical cues on cancer cell invasion and metastasis.

## Introduction

Metastasis is the process of cancer cells disseminating from primary or metastatic tumors, followed by the invasion of other organs or tissues and the subsequent formation of secondary tumor sites.^1^ The first key event in the metastasis cascade is the infiltration of single cells or cell clusters into the surrounding extracellular matrix (ECM).^2, 3^ As this invasion is heavily influenced by the tumor microenvironment, research has concentrated on understanding the local mechanisms driving the metastatic transformation with the aim to find strategies for inhibition.^4^ While biochemical traits have been a long-standing focus of analysis, mechanical cues are increasingly considered a critical contributor to malignant cancer transformation and metastasis. The mechanical properties of the tumor and individual cells, however, are not the only factors affecting metastasis. The surrounding tumor microenvironment also provides essential biophysical cues such as matrix stiffness and tensile forces that critically modulate the metastatic process.^5–7^ These and other local physical properties are affected by the dynamic relationship between cancer cells and their surrounding microenvironment, which is characterized by alteration of tissue composition and architecture in a manner that supports malignant transformation and metastasis.^6, 8, 9^ The tumor stroma plays here a key role as it is populated by stromal cells such as fibroblasts and immune cells. Fibroblasts in particular have been shown to dynamically remodel the tissue microenvironment by exerting contractile forces of several tens of nN within the surrounding ECM.^10^ Furthermore, both cancer cells and fibroblasts, can cause strain fields in their surrounding matrix which can well exceed 100 μm in diameter.^10, 11^

To study the effect of biophysical forces on malignant progression, cancer models are increasingly orienting towards three-dimensional (3D) culture matrices and away from historic approaches of two-dimensional (2D) cell culture performed on rigid plastic substrates. Composed of synthetic or biopolymers, these 3D models more accurately recapitulate the ECM environment and can be designed with specific stiffnesses, architectures, and the presence of surface ligands.^12, 13^ Moreover, considerable progress has been made in the reconstitution of biomimetic cell culture models, such as through advances in 3D tumor models, implementation of engineered models for the surrounding tumor microenvironment, and the reorganization of *in vitro* models from patient-derived samples for personalized medicine approaches.^14, 15^ Utilizing these 3D matrices to investigate cancer cell invasion, and the interaction of individual cancer cells or cell clusters with the surrounding matrix is an important step for physiologically relevant studies. For instance, studies conducted in a 3D *in vitro* environment have demonstrated the significance of matrix stiffness on cancer cell migration and matrix invasion, with single-cell invasiveness exhibiting a biphasic response to increasing Collagen I concentrations.^16, 17^ While such models have greatly increased the physiological relevance of applied cell culture systems, the high number of resistances against chemotherapeutic responses suggests that *ex vivo* model systems might profit from going beyond solely adapting the composition and architecture of in vitro tissue models.^18^ The active role of tumor-associated cells in the deformation of living tissue suggests that triggering a dynamic deformation to the surrounding tumor matrix can critically contribute to understanding the microenvironment experienced by cancer cells. To investigate the effect of matrix deformation on cancer cell migration, different approaches have been implemented.^19, 20^ A particularly interesting approach is the application of remotely controlled microscale actuation of *in vitro* systems. One promising strategy is magnetically controlled micro- or nanoscale actuators embedded in samples with the capability to induce microscale deformations of the ECM. The remote control offers a non-invasive method and is compatible with sterile cell culture environments. A study performed by Menon *et al.,* employed microparticles embedded in a 3D matrix, which were actuated using a permanent magnet to achieve a transient tugging force that was shown to enhance cancer cell invasion into 3D collagen matrices in a fibronectin-dependent manner.^21^ The same research group reported a cell-line-specific response to different types of mechanical stimuli such as matrix deformation and rigidity. They used a combination of substrates with varying degrees of compliance and triggered a cyclic deformation of the matrix using a single magnetic bead.^22^ Aiming to test the effect of local deformation of the ECM on cells residing there, further approaches have been developed but are often focused on individual cells and are not compatible with established 3D cancer invasion assays.^23^ In a recent study by Shou *et al.*, 3D magnetic hydrogels were developed to study the effect of stiffness changes on embedded cancer spheroids. Their findings revealed that matrix stiffening promoted malignancies, while matrix softening downregulated the expression of mechano-transductors.^24^ While, in that study, the whole matrix stiffness could be rapidly altered to measure downstream effects, the influence of dynamic mechanical compression and strain of the network generated from individual point sources, like individual cells, remains to be studied.

This work applies magnetically controlled microscale actuation of 3D tumor invasion assays over several days of culture. Surface functionalized anisotropic rod-shaped iron microparticles (μRods) were covalently linked to a fibrous 3D Collagen I (Col I) network, which was employed as an ECM model in cancer spheroid invasion assays. Magnetically controlled cyclic actuation of the sample-embedded μRods in the vicinity of the tumor spheroids allowed for local deformation of the tumor microenvironment. Using particle velocimetry analysis, we determined the range of Col I deformation in response to the magnetic field which spanned several times the length of a single μRod—values comparable with strain fields reported to originate from contracting cells.^10^ Moreover, the patterns of Col I deformation resemble findings published for cells residing in Col I hydrogels.^10, 11^ Changes in cellular calcium levels were observed in response to sustained actuation of the cell-embedding Col I matrix, indicating that calcium signaling might be influenced by the mechanical dynamics of the cellular microenvironment. Sustained actuation of the ECM model surrounding spheroids of the invasive cancer cell line MDA-MB-231 at 1 Hz over two days resulted in increased dissemination and invasion of cancer cells into the surrounding Col I network. Locally concentrated application of the μRods in the vicinity of Col I embedded spheroids was shown to cause local attraction of the adjacent cancer cells and patterns of higher cell densities in the direction of the applied mechanical stimuli. Overall, this study suggests that local micromechanical deformation can promote cancer cell invasion and provides means to apply and test the influence of controllable non-invasive micromechanical cues in 3D *in vitro* systems.

## Experimental Section

### Cell Culture

GFP labelled cells from the invasive cancer cell line MDA-MB-231 were ordered from GenTarget Inc. Cells were cultured in complete medium (Gibco DMEM, high Glucose, GlutaMAX Supplement, pyruvate, Fisher Scientific) supplemented with 10 % fetal bovine serum (FBS, Biowest) and 1% penicillin/streptomycin (P/S, 10 000 u/mL, Thermo Scientific) at 37°C and 5% CO2. Cells were passaged at 70%-80% confluency by washing cells once with Dulbecco’s phosphate buffered saline (Gibco DPBS, no calcium, no magnesium, Thermo Fisher Scientific), incubation with Trypsin-EDTA (0.25%, phenol red Thermo Fisher Scientific) and subsequent neutralization of the trypsin using complete medium. Cells were then concentrated by centrifugation, counted, and used for experiments or directly reseeded for further culture.

### Col I Hydrogel Preparation

Col I hydrogels were prepared as described earlier.^25^ Briefly, Col I hydrogels were prepared from high-concentration rat-tail Col I (Corning, #354249, 9.48 mg/ml) following the manufacturer’s instructions. Concentrated Col I was mixed with cell culture medium and 0.5 M NaOH to adjust the mix to pH 7 to allow for polymerization. The sample was thoroughly mixed and incubated at 37°C for a minimum of 30 minutes. For the preparation of fluorescently labelled Col I hydrogels, 10% of the Col I stock was replaced by fluorescently labelled Col I stock, prepared according to the protocol specified in the following section.

### Fluorescent Labeling of Col I Hydrogels

Fluorescent labelling of Col I was performed according to a previously published protocol.^25^ Briefly, a stock solution of 5(6)-TAMRA N-succinimidyl ester (C0027, Chemodex) was prepared in dimethyl sulfoxide (DMSO) at a concentration of 10 mg/mL, vortexed thoroughly and stored under exclusion of light at −20°C. 2 mL of concentrated Col I was injected to a 3 mL 3.5k MWCO Slide-a-Lyzer Dialysis Cassette (Thermo Fisher) and dialyzed overnight against 2 L of sterile filtered aqueous labeling buffer (0.25 M NaHCO3, 0.4 M NaCl, pH 9.5) at 4°C. All subsequent steps were performed under exclusion of light.

The dialysis cassette was removed from the labeling buffer, 100 μL of TAMRA stock solution was mixed with 900 μL of labeling buffer and injected to the dialysis cassette. The cassette was wrapped in aluminum foil and incubated on a rocking shaker overnight at 4°C. Next, to remove excess dye and reconstitute the acidic pH of the Col I stock, the dialysis cassette was thoroughly washed in acidic washing buffers. First, the cassette was placed in 1 L of 0.5 M acetic acid at pH 4 and dialyzed at 4 °C overnight. Next, the cassette was subjected to two subsequent dialysis washes in each 1 L of 0.02 M acetic acid at pH4 overnight at 4°C. The fluorescently labelled Col I was withdrawn from the dialysis cassette and stored at 4°C under exclusion of light.

### Fabrication, functionalization and characterization of magnetic iron μRods

Magnetic iron μRods were fabricated using template restricted electrodeposition. The method to prepare plating templates via photolithography was developed orienting on work published by B. Özkale.^26^ Further, the Magnetic iron μRods are functionalized and characterized, see details in **Supplementary Text 1** and **Supplementary Figures S1-S4**.

### PDMS Pool Preparation

PDMS pools were prepared as previously published.^25^ Briefly, Polydimethylsiloxane (PDMS) polymer (SYLGARDTM 184 Silicone Elastomer kit, Dow Corning #01673921) was mixed according to manufacturer’s instructions and degassed for 1 hour. The degassed PDMS mix was poured onto a blank wafer to obtain a layer thickness of 4 mm, degassed again for 1 hour and cured at 80°C for 3 hours or overnight. The cured PDMS disc was detached from the wafer and rings were punched out with an inner diameter of 8 mm. Using oxygen plasma treatment, PDMS pools were bonded to glass cover slips and sterilized by washing with 70% ethanol in deionized water and UV treatment for a minimum of 30 minutes.

### Surface Functionalization of Glass and Plastic Surfaces

To enhance adhesion of Col I hydrogels to commercially available and self-fabricated sample substrates made from glass, PDMS or plastic, a surface modification with dopamine was performed, according to a protocol published by Park et al.^27^ If not already delivered in sterile packaging, the sample to be surface-treated was sterilized by thorough washing using 70% Ethanol in deionized water and subsequent exposure to UV-radiation for at least 30 minutes. Next, the surfaces of interest were incubated with a sterile filtered solution of 2 mg/mL dopamine (Dopamine 62-31-7, Sigma Aldrich) in 10 mM Tris-HCl (pH 8.5) at room temperature for 2 hours. During the incubation, a change of color from transparent to dark grey-brown could be observed, indicating the progression of pH-induced oxidation. After incubation, the reaction solution was aspirated, samples were gently rinsed with autoclaved deionized water and left to dry. Until further use, samples were stored under sterile conditions in sealed plastic dishes at 4°C.

### Magnetic Actuation Using an Electromagnetic Field Generator

To infer suitable magnetic field parameters and allow live confocal imaging during magnetic actuation, an eight-coil electromagnetic field generator (MFG 100-I, Magnebotix AG) was used that was mounted on a spinning disk confocal microscope. Field magnitudes exceeding 20 mT were achieved using conical field focussing core extensions, resulting in a spherical workspace of 1 mm radius. Deflections upon actuation were analyzed optically for sweeps of field magnitudes and frequencies.

### Long-term Magnetic Actuation Using a Custom-built Halbach Array

For cyclic magnetic actuation of multiple samples in parallel over several days, an incubator-compatible cylindrical Halbach array was built (**Supplementary Figure S5**). This Halbach array comprised eight positions of permanent magnets (NdFeB, N42) with dimensions of 0.5×0.5×2.0 cm arranged in a circular array. To obtain a cylinder of 8 cm in length, four of the bar magnets were placed in series at the magnet positions along the longitudinal axis of the cylindrical array. The dimensions of the cylindrical magnet array were determined using Finite Element Method Magnetics (FEMM) ^28^ and designed to result in a uniform magnetic field of 50 mT within a cylindrical workspace of 3 cm. The Halbach aray actuator was 3D printed from Polyethylene terephthalate glycol (PETG) using a filament extruder and assembled with a motor (P25UK12, PiBorg) and a speed controller (Global Specialities) inside a custom-designed acrylic casing (0.5 cm thickness). The components were stably connected using dichloromethane (Sigma-Aldrich) and protected from humidity using silicon paste and thermoresistant tape. The rotational frequency of the assembled setup was measured and adjusted using a hand-held tachometer (PeakTech 2790) to 1 Hz. The magnetic field of approximately 50 mT was confirmed inside the Halbach cylinder using a Hall Probe (Metrolab 3D Magnetometer) that was positioned using a piezo controller mounted on a custom-made 3D printed holder. Measurements of various positions (performed in triplicate) are displayed in **Supplementary Figure S6**.

### Fluorescence and Brightfield Imaging

Fluorescence imaging was performed using a Nikon Eclipse Ti2 microscope that was equipped with ta CSU-W1 Confocal Scanner Unit and a Hamamatsu C13440-20CU ORCA Flash 4.0 V3 Digital CMOS camera. The Nikon NIS-Elements Advanced Research 5.02 (Build 1266) software was used for operation of the microscopy and image acquisition. Images were acquired with Nikon CFI Apochromat objectives (10x, 20x, 40x). See **Supplementary Text 2** for image processing analysis.

### Collagen Deformation with Strain Rate Mapping

For Col I deformation studies, 700 μL of Col I hydrogel were prepared at a final concentration of 2 mg/mL as specified earlier. During preparation, 10 µL of the complete medium were substituted with a suspension of fluorescent microparticles (Fluoromax, 0.2 μm, R200, Thermo Scientific) prior to addition of the remaining components. Collagen hydrogels were transferred to PDMS wells with 4 mm diameter and left to polymerize for 45 minutes at 37°C, 5% CO2 in a humidified petri dish. To track collagen deformation, actuation experiments were conducted during live imaging using the magnetic field generator^29^ mentioned above. Actuation was performed at 50 mT field magnitude with fields rotating at a frequency of 1 Hz either in in-plane (xy), or out-of-plane (xz). Video sequences of 30 seconds were acquired with a 40x objective using confocal mode imaging in the TEXAS red channel (580 nm), see **Supplementary Videos V1** and **V2**.

Col I hydrogel deformation was recorded at a frame rate of 100 ms and the displacement of the fluorescent tracer particles was analysed via Particle Image Velocimetry using the PIVlab plugin for MATLAB.^30–33^ Image sequences were preprocessed by enabling the CLAHE (Contrast Limited Adaptive Histogram Equalization) function. PIV analysis was then performed using the FFT window deformation algorithm with an initial interrogation area of 128 pixels followed by two additional passes with decreased window size (halving the pixel number in every consecutive pass). The vectors indicating local displacement in m/s were scaled to 10 and superimposed with a display of the simple strain rate [1/s] presented in a color map. As the simple strain rate depicts the velocity gradient relative to the local velocity, their value indicates the direction of particle displacement, resulting in both positive and negative values within the obtained set of data.^34^

### Ca^2+^ Imaging Experiments

3D Col I matrices that were populated with MDA-MB-231 cancer cells and enriched with surface-functionalized iron μRods for Ca^2+^ imaging in the presence and absence of magnetic actuation. 50 μL of Col I at a concentration of 2 mg/mL were added per PDMS well and incubated in a humidified chamber at 37°C. Next, MDA-MB-231 cells were dispersed in 2 mg/mL Col I matrices at a concentration of 10^5^ cells/ mL, 5 μL of iron μRods were added per 200 μL of Col I-cell mixture and 70 μL of the cell-Col I mixture were applied on the samples. Samples were incubated for 1 h, then covered with cell culture medium and further incubated for 3 days at standard incubator conditions. 1.5 h prior to the experiment, Col I samples were once gently washed with PBS and incubated with 10 μM solution of Calbryte 520 AM (AAT Bioquest), diluted in a labeling buffer prepared from Modified Hanks Buffer with Calcium and Magnesium, supplemented with 0.04% Pluronic F-127 and 20 mM HEPES. Samples were incubated at 37°C, 5% CO2 for 1 h and gently washed twice with the labeling buffer prior to imaging.

Live imaging was performed while samples were actuated using the magnetic field generator described earlier (**Supplementary Video V3**). During the experiment, image sequences were acquired using the GFP channel with a frame rate of 200 ms to capture the Ca^2+^ influx while the deflection of the iron μRods was inferred separately via the brightfield channel. Images were analyzed using FIJI and the Raw Integrated Density was measured in regions of interest (ROI), representing the overall signal intensity within the selected ROI.

### Spheroid Preparation

Spheroids were prepared using low adhesion round bottom 96 well plates (Corning, Costar Ultra-Low Attachment 96-well plate). 5×10^3^ cells were seeded per well in 100 μL of complete medium and plates were centrifuged at 500 g for 5 minutes. After one day of culture, 100 μL of media containing 10 μg/mL Col I were added per well to obtain a final concentration of 5 μg/mL. The plate was centrifuged again at 500 g for five minutes and incubated for two more days.

### Spheroid Embedding

Spheroids were embedded in Col I hydrogels after three days of culture. To enhance adhesion of Col I hydrogels, 8-well culture slides (μ-Slide 8 well, ibiTreat, IBIDI) were surface functionalized with dopamine as described above and briefly incubated with complete medium prior to use. 150 μL of Col I hydrogels prepared at the desired concentration were added per well. Spheroids were collected with a 200 μL pipette and transferred to Col I hydrogels, releasing them gently from the pipette without disruption of the collagen network and aiming to keep the amount of released medium to a minimum. Col I hydrogels were left to polymerize for approximately 30 minutes in a cell culture incubator at 37°C and 5% CO2. Next, an additional layer of 100 µL of Col I was added on top of the previously prepared samples, either enriched with functionalized μRods or not. Samples were left to polymerize for 30 more minutes at 37 °C. To prevent dehydration of the hydrogels, samples were incubated in humidified chambers within the cell culture incubator.

## Results

### Analysis of Tumor Cell Invasion into 3D Col I Hydrogels

Cancer cell invasion marks the first step of the metastatic cascade (**Figure 1a**). To model cancer cell invasion *in vitro*, 3D tumor spheroids from the invasive breast cancer cell line MDA-MB-231 were embedded in 3D Collagen I (Col I) matrices and imaged over three days (**Figure 1b**). As previously reported, our results confirmed that the invasion of tumor cells into the surrounding matrix correlates with Col I concentration in a biphasic manner.^35^ Cancer cell invasion correlated with increasing concentration of Col I hydrogels between 0.5 mg/mL to 3.0 mg/mL and a peak of invasive behavior was identified for Col I matrices at a concentration of 3 mg/mL. Higher concentrations of Col I hydrogel negatively correlated with cancer cell invasion (**Supplementary Figure S7**).

**Figure 1:**
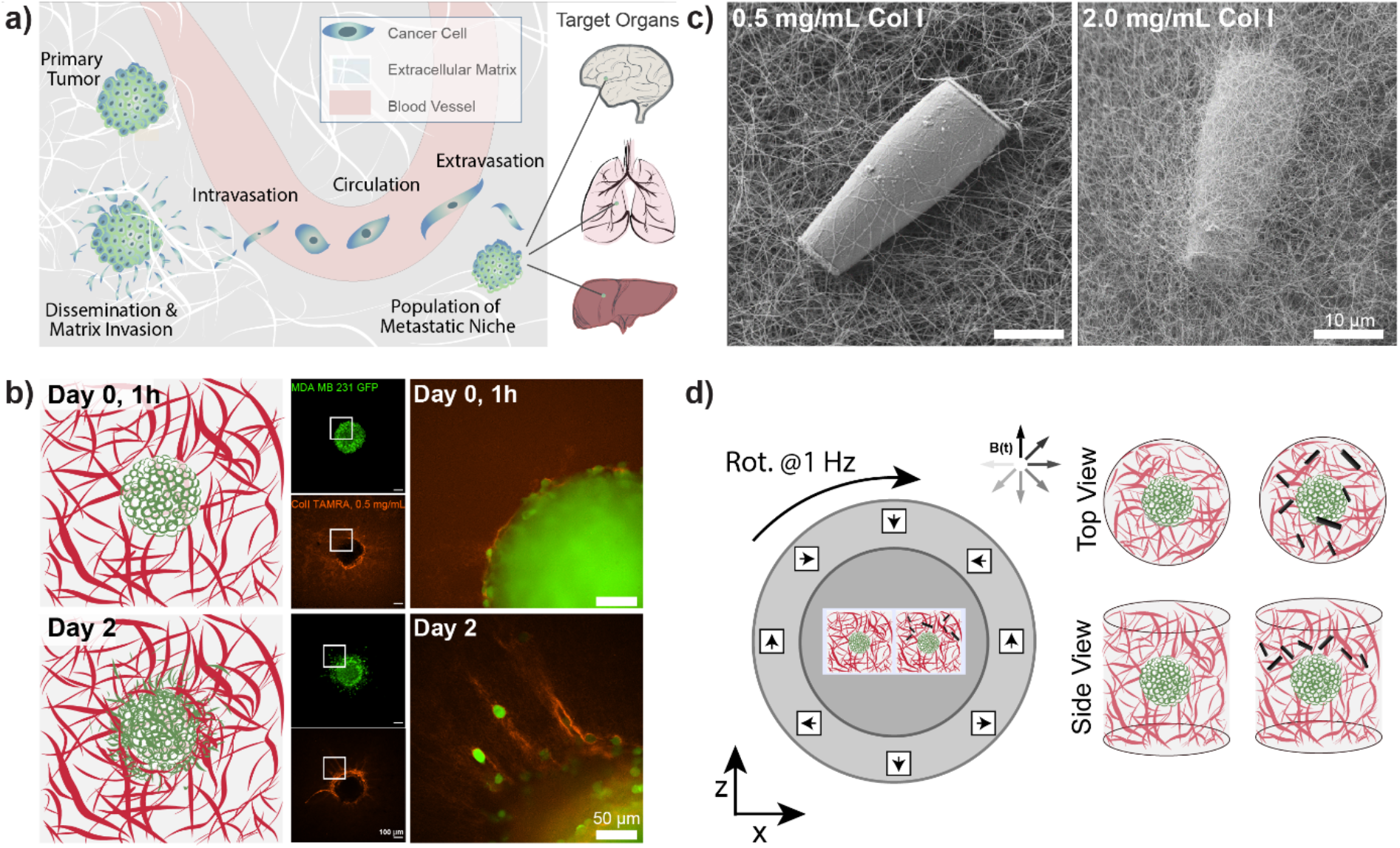
Magnetically controlled deformation of in vitro cancer environments. a) The metastatic cascade is initiated by dissemination of cancer cells from primary tumors followed by the invasion of surrounding tissues. From there, a fraction of invasive cancer cells successfully reach and enters blood vessel, travel with the blood stream and eventually enter distant tissues or organs where they form secondary tumor sites. The brain, lungs and the liver are among the organs which are most frequently affected by metastasis. b) To study cancer invasion *in vitro*, tumor spheroids of the invasive breast cancer cell line MDA-MB-231 (shown in green) were embedded in a Col I matrix (0.5 mg/mL concentration, fluorescently labelled in red) and cancer cell invasion was repeatedly monitored over several days of culture. Scale bars: 50 μm. c) Scanning electron micrograph of rod-shaped magnetic microparticles (μRods) embedded in a Col I matrix of 0.5 mg/mL (left) and 2.0 mg/mL (right) concentration. d) Schematic display of the experimental setup to study cancer invasion during magnetically controlled deformation of the extracellular matrix over several days of culture. A uniform, rotating magnetic field of 50 mT was generated using a cylindrical Halbach and was employed to actuate magnetic μRods that were embedded in 3D tumor invasion models.

Fluorescent labelling of Col I hydrogels used for cancer invasion studies showed densification and remodeling of the Col I hydrogel network surrounding the invading tumor cells. Stronger patterns of network deformation were observed at lower concentrations of Col I hydrogels, potentially caused by embedded cells and enhanced optical resolution of individual Col I. Representative examples of tumor spheroids cultured in different concentrations of fluorescently labelled Col I matrices are displayed in **Supplementary Figure S7b-d**.

For the following studies on the influence of cyclic deformation of Col I hydrogel matrices on the invasive behavior of embedded tumor spheroids, a Col I hydrogel concentration of 2 mg/mL was used. While a concentration of 3 mg/mL was found to result in the most pronounced spreading of cancer cells to the surrounding matrix (as displayed in **Supplementary Figure S7b**), a concentration of 2 mg/mL showed similar behavior of matrix invasion while still not reaching maximum values. Hence, while allowing for the reconstitution of an invasive phenotype, a system was selected which did not maximize invasiveness. This way, changes in invasiveness in response to local ECM deformation could be observed relative to the negative control condition without actuation.

### Magnetically Controlled Actuation of *In Vitro* ECM Models

Tumor cells interact and invade their surrounding microenvironment, which is also populated by other cell types, such as highly contractile fibroblasts. **Figure 1b** illustrates how individual cancer cells disseminate from their original location (here a tumor spheroid) and invade the surrounding Col I matrix while further reorganizing the surrounding fiber structure. Our study aimed to investigate how cancer cell invasion could be influenced by cyclic local deformation of the tumor microenvironment, mimicking the interaction of cells residing in the tissue with their surrounding ECM network.

Magnetic μRods, approximately 25 µm in length and 3.5 µm in radius, covalently linked to the Col I hydrogels (**Figure 1c**) were used in 3D tumor invasion models as ECM-embedded microscale mechanical actuators. Acute cytotoxicity of the iron μRods was excluded by an MTT assay, as displayed in **Supplementary Figure S4**. To determine the field magnitude used for the actuation experiments, Col I hydrogel samples were prepared at a concentration of 2 mg/mL and mixed with surface-functionalized μRods. Using a versatile electromagnetic field generator mounted on a confocal fluorescence microscope, rotating magnetic fields (RMF) of varying magnitude were tested to determine a field magnitude that would allow for readily detectable deflection of the Col I -embedded μRods without disrupting the surrounding Col I network. The experimental setup is depicted in **Supplementary Figure S8**. A working RMF magnitude of 50 mT was determined, and hereafter a RMF of 50 mT with 1 Hz frequency was adopted for collagen defection and cell invasion studies. In addition, to enable exposure of multiple tumor spheroid samples to a uniform RMF over several days under standard cell culture conditions, a custom incubator-compatible sample actuation system based on a rotating Halbach cylinder was built. The device consists of permanent magnets arranged in a first order Halbach cylinder^36^ (**Figure 1d**) designed to produce a magnitude of 50 mT. It is mounted laterally on a ball bearing and coupled to a motor that drove its rotation around the x-axis with a frequency of 1 Hz.

### Characterization of Magnetically Controlled Col I Hydrogel Deformation

Live imaging of the deformation of Col I hydrogels through magnetic actuation of μRods embedded in samples was investigated using the microscope-mounted electromagnetic field generator. In all experiments, the same actuation parameters previously determined and described for the incubator-compatible Halbach actuation setup (50 mT, 1 Hz) were applied. To visualize Col I hydrogel deformation during magnetic actuation, hydrogel samples were supplemented with fluorescent polystyrene nanoparticles (200 nm diameter) and imaged via confocal fluorescence microscopy. The resulting deformation patterns of the Col I matrices by the embedded magnetic μRods were analyzed for both in-plane rotation (about the z-axis) and out-of-plane rotation (about the x axis) under the specified RMF conditions. We then quantified the strain rates resulting from the cyclic deflection. Results for in-plane and out-of-plane RMF are displayed in **Figure 2**, and **Figure 3** respectively. In both modes of actuation, the extent of the strain rates was found to change over the course of one cycle, as visualized by the change in color intensity in the overview maps of strain rates plotted over time in **Figure 2** Error! Reference source not found. and **Figure 3**. The torque applied in both modes was estimated to be on the order of 80 pNm for the rods, with details of the torque calculation provided in **Supplementary Text 3**. This magnitude is similar to previously reported cell contractile moments exhibited e.g. by fibroblasts.^37–40^

**Figure 2:**
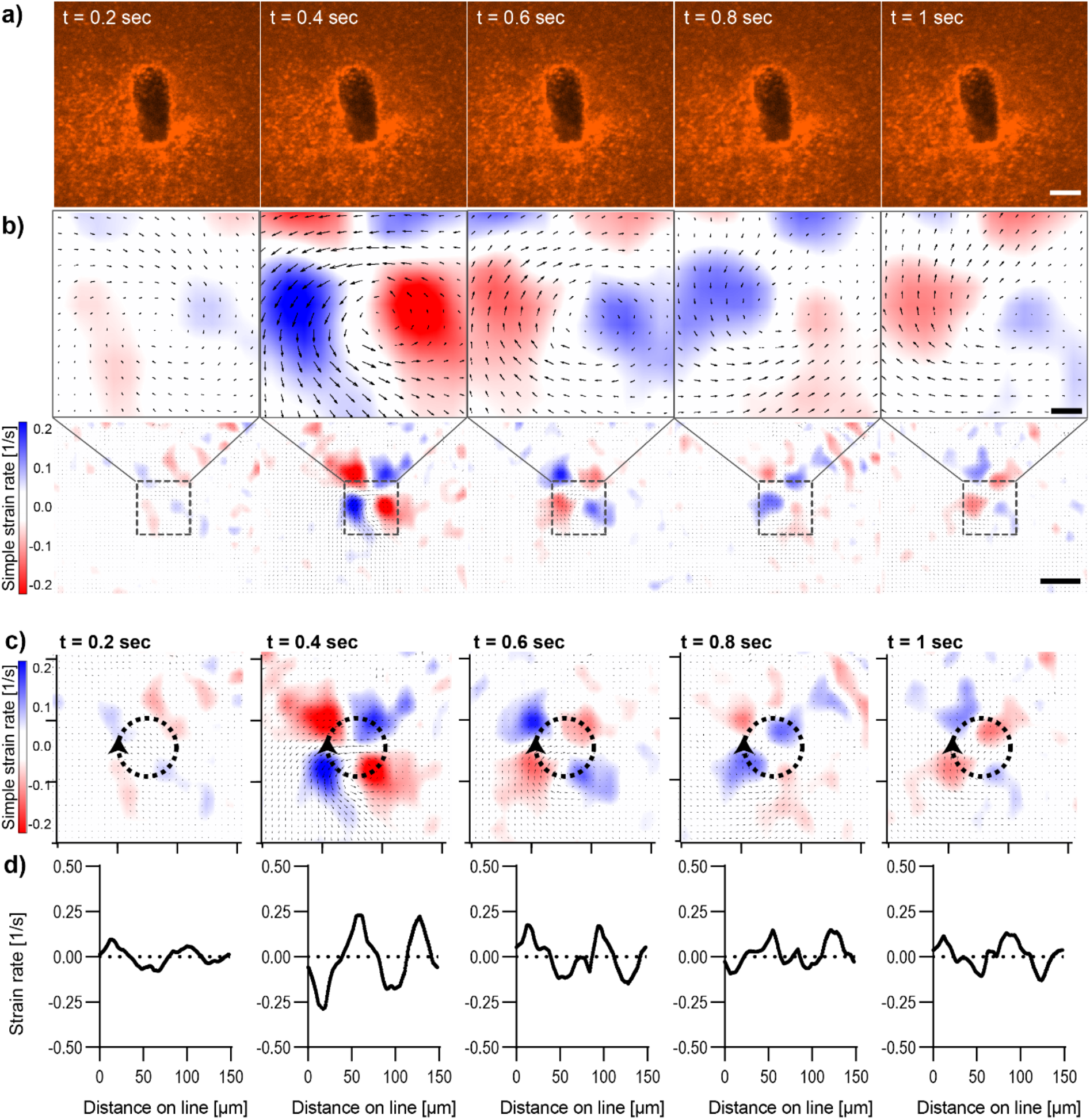
Analysis of Col I hydrogel deformation upon in-plane deflection of an embedded μRod. Magnetic actuation was performed using an in-plane rotating magnetic field of 50 mT magnitude with a rotational frequency of 1 Hz. Images were acquired with a frame rate of 100 ms. a) Confocal fluorescence micrographs of Col I hydrogels (2 mg/mL) enriched with fluorescent tracer beads (red) over the course of one actuation cycle. Scale bar: 10 μm. b) Particle Image Velocimetry analysis of the image sequence depicted in a). The top row shows the equivalent image regions as displayed in row a), analyzed for the simple strain rate extracted for different timepoints. The bottom row of section b) displays the image areas in the context of a larger field of view. Scale bars top row: 10 μm, bottom row: 50 μm. c) Simple strain rates were extracted along the circumference of circular regions surrounding a deflecting μRod. Ticks of orientation axes indicate 50 μm. d) Plot of data extracted from the strain rate mapped in c), the arrowhead marks the onset and direction of the line along which strain rates were extracted from the map.

**Figure 3:**
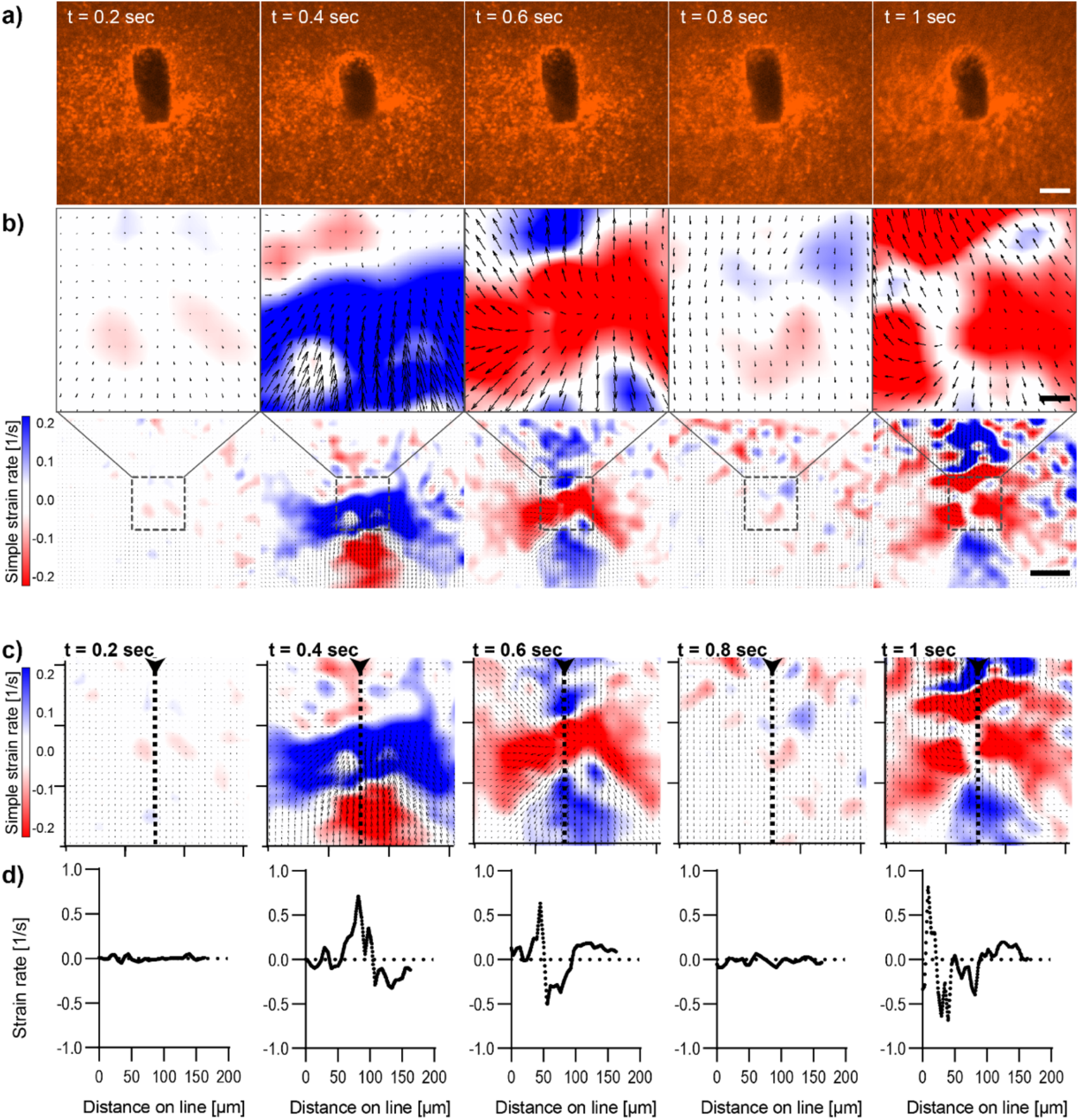
Analysis of Col I hydrogel deformation upon out-of-plane deflection of an embedded μRod. Magnetic actuation was performed using an out-of-plane rotating magnetic field of 50 mT magnitude with a rotational frequency of 1 Hz. Images were acquired with a frame rate of 100 ms. a) Confocal fluorescence micrographs of Col I hydrogels (2 mg/mL) enriched with fluorescent tracer beads (red) over the course of one actuation cycle. Scale bar: 10 μm. b) Particle Image Velocimetry analysis of the image sequence depicted in a). The top row shows the equivalent image regions as displayed in row a), analyzed for the simple strain rate extracted for different timepoints. The bottom row of section b) displays the image areas in the context of a larger field of view. Scale bars top row: 10 μm, bottom row: 50 μm. c) Simple strain rates were extracted along the axis of deflection of the actuated μRod. Ticks of orientation axes indicate 50 μm. d) Plot of data extracted from the strain rate mapped along the indicated dashed line shown in c), the arrowhead marks the onset and direction of the line along which strain rates were extracted from the map.

In-plane rotation of the magnetic field was shown to cause a radially symmetric pattern of strain rate distribution around the center of rotation of the magnetic μRod (**Figure 2b-d, Supplementary Movie V1)**. Maximal values were found at 0.4 s after the onset of deflection and reached values of simple strain rates of approximately ±0.2 /s The extraction of strain rates along the circumference of a circle surrounding the center of μRod deflection with an assigned radius of 24 μm further displays the pattern of alternating positive and negative values of strain rates surrounding the center of μRod deflection in four quadrants (**Figure 2b-c**). This behavior is correlated with the direction of Col I network displacement during the rotation of the actuated μRod. Considerable strain is detectable within regions spanning 200 µm in diameter around the µRod, which is in line with reported strain fields generated by cells, as noted earlier.^10, 11^

The analysis of image sequences acquired during out-of-plane actuation of magnetic μRods within the Col I hydrogel was performed analogously to the in-plane magnetic field rotation and is shown in **Figure 3 (Supplementary Movie V2)**. Here, as expected for any cyclic deformation, two directions of Col I displacement were detected. However, the strain rates measured during out-of-plane rotation were more than double the strain rate values observed for in-plane field rotations. The pattern of Col I deformation caused by out-of-plane defection of a μRod displays regions expressing positive and negative strain rates, respectively, to extend laterally, perpendicular to the long axis of the μRod, hence along the axis of the field rotation (**Figure 3b-c**). The values of the strain rates identified in the analysis are shown to vary in a periodic pattern, representative values from one cycle of field rotation are shown in the image sequences of **Figure 3**. In an originally relaxed state (**Figure 3b**, 0.2 s) strain rates are low, reflecting only marginal particle displacement. During the course of deflection, strain rates are increasing, as shown at timepoints t= 0.4 and t= 0.6 s, respectively, resulting from a larger displacement of tracer particles per time, and thus, increased particle velocity. At t=0.8 s, a planar position of the μRod within the image plane is again observed, similar to the fluorescence micrograph shown for timepoint t=0.2s.

The values extracted from the region of the center of deformation (**Figure 3c-d**) report on the locally identified strain rates that start close to zero and assume peak values of approximately 0.7 /s (for positive strain rates) and close to −1 /s for the negative strains rates, as observed at timepoint t = 1 s of the magnetic field rotation. The extraction of strain rates along a line perpendicular to the axis of rotation confirms the alternation of positive strain and negative strain rates within the area depicted in the strain rate maps (**Figure c-d**). Furthermore, the strain rates extracted from image sequences during the actuation of a μRod caused by the out-of-plane rotation of a magnetic field show detectable strains developed over an area exceeding 200 μm in diameter, as indicated in the overview maps in **Figure 3b**, bottom. The higher values may be explained by the stronger volumetric contribution of underlying planes of the network which are affected during this actuation mode on contrast to in-plane rotation, given that the collagen embedded µrods are added as another layer atop. In summary, both modes provide means to generate physiologically relevant strain fields originating from individual localized sources of single cell dimensions.

### Magnetically induced mechanical deformation triggers increased Calcium influx

Having quantified the generation of strain fields and confirmed their relevant matrix deformation, we next sought to test their impact on cancer cells. Stretch sensitive ion channels are a prime example for mechanically triggered signaling processes in living cells. Embedded in the cellular membrane, they are activated by deformation of the membrane and allow for the influx of calcium ions that trigger intracellular signaling processes. To test whether the mechanical forces resulting from magnetic actuation of Col I-embedded iron μRods provoke a change in Ca^2+^ levels within living cells, experiments were performed to investigate potential changes in cellular Ca^2+^ levels during actuation of Col I networks surrounding MDA-MB-231 cancer cells. Cancer cells were embedded with µRods in Col I hydrogels (at a concentration of 2 mg/mL). By means of a fluorescent Ca^2+^ indicator probe, changes in intracellular Ca^2+^ levels were optically monitored during magnetic actuation of the embedded μRods (**Figure 4**). Image sequences recorded of cell samples treated with the fluorescent Ca^2+^ indicator probe were analyzed for fluorescence intensity levels prior to, during, and after cyclic magnetic actuation of the μRods embedded in the vicinity of Col I-embedded cancer cells. (**Figure 4a**). Prior to actuation (**Figure 4a, i) and 4b**, t=0 sec), images sequences of 1 minute show a low and comparably homogenous fluorescence signal. Upon actuation of the sample with a magnetic field of 50 mT at 1 Hz frequency for 1 minute, the repeated exposure of the cells to a detectably deflected μRod (as depicted in **Figure 4a, ii**)) is correlated with an increase of fluorescence intensity in the region of μRod-adjacent cancer cells (see also **Supplementary Video V3**). This trend is detected as prominent prominent peak in the fluorescence values that are plotted as function of time in **Figure 4c**. Correlating fluorescence intensity with levels of intracellular Ca^2+^, our data suggests that repeated deformation of the cellular environment by adjacent μRods triggers the influx of Ca^2+^ ions into the cellular body.

**Figure 4:**
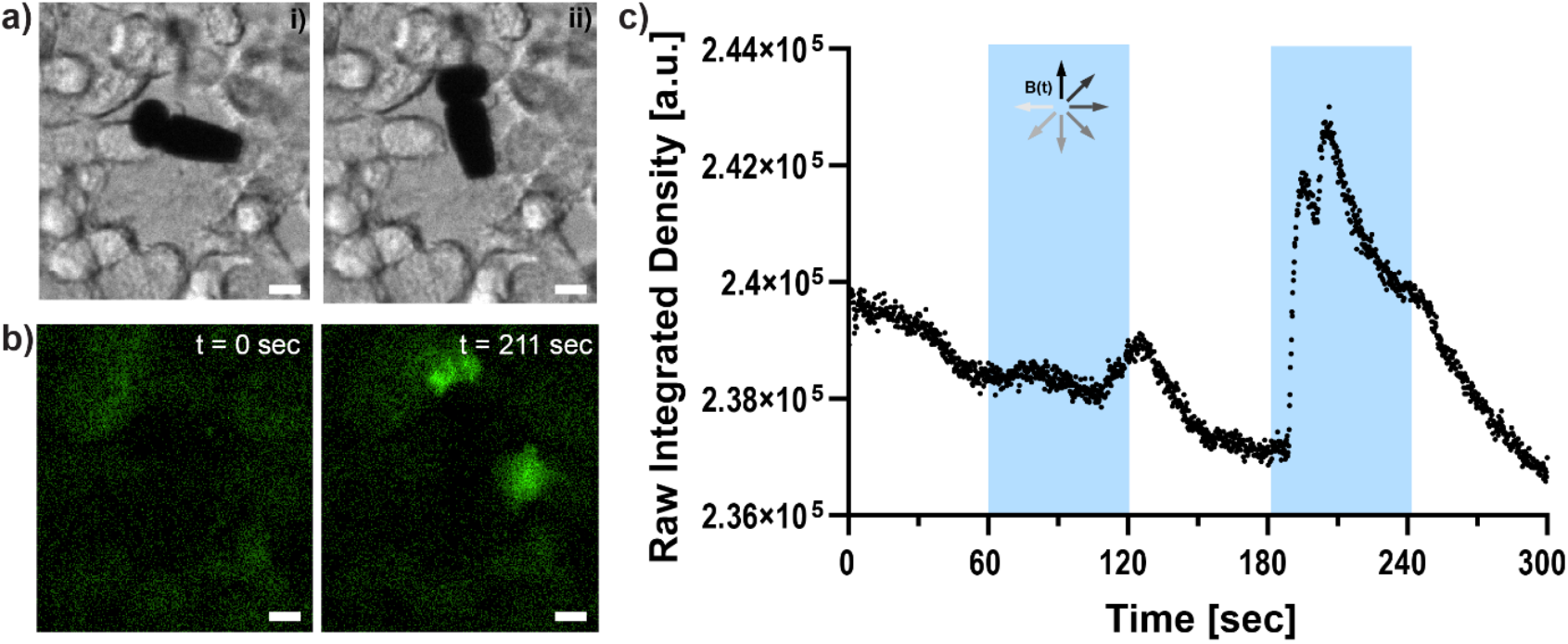
Actuation of hydrogel-embedded μRods results in signals indicating Calcium influx to hydrogel-residing MDA-MB-231 cells. a) Bright field microscope image of a functionalized μRod embedded within a Col I hydrogel matrix populated with MDA-MB-231 cells. Application of an in-plane rotating magnetic field of 50 mT results in deflection of the μRod within the hydrogel matrix. Different orientations of the μRods resulting from magnetic field orientation are shown in i) and ii). MDA-MB-231 cancer cells were stained with a Calcium indicator dye and imaged before and during exposure to an in-plane rotating magnetic field of 50 mT. Images show cells in the vicinity of μRods before (left, t=0 s) and after actuation of the samples (t=211 s chosen as representative timepoint). All scale bars: 10 μm. c) Fluorescence intensity was analyzed during cycles of magnetic field actuation and is shown as Raw Integrated Density. Blue shading indicates timespans of sample exposure by rotating magnetic field.

### Micromechanical deformation of Col I hydrogels increases tumor cell invasion

Next, the influence of long-term cyclic micromechanical deformation of the tumor microenvironment on the invasive behavior of MDA-MB-231 spheroids was investigated. Based on the previously described tumor invasion model, spheroids prepared from MDA-MB-231 GFP cells were embedded in Col I hydrogels of 2 mg/mL concentration. A second layer Col I hydrogel containing the surface functionalized iron μRods was then applied. Imaging was performed 1 h post spheroid embedding, before samples were exposed to RMF. Repeated acquisition of fluorescence data was performed after 18 h and 42 h of incubation with or without magnetic actuation, respectively (**Figure 5a**). As displayed in **Figure 5b**, invasion of tumor cells into the surrounding Col I hydrogel matrix was significantly increased after 42 h of magnetic actuation compared to the control condition, where spheroids were cultured in the presence of μRods but without exposure to a rotating magnetic field. Furthermore, the invasive behavior of MDA-MB-231 cells was neither expected to be appreciably affected by the application of RMF alone, nor did our experiment reveal a statistically significant effect (95% confidence interval) on samples without µRods that were exposed to RMF, as compared to unexposed control samples. Representative images of the spheroids imaged before and after incubation in the presence and absence of a RMF and μRods are displayed in **Figure 5c** (before incubation) and **Figure 5d** (after 42 h of incubation). The yellow outline indicates the boundary that was defined for the quantification of spheroid spreading. As apparent from the data obtained from the Feret diameter calculations displayed in **Figure 5b**, the spread of tumor cells into the surrounding Col I matrix is readily detectable in the fluorescence images.

**Figure 5:**
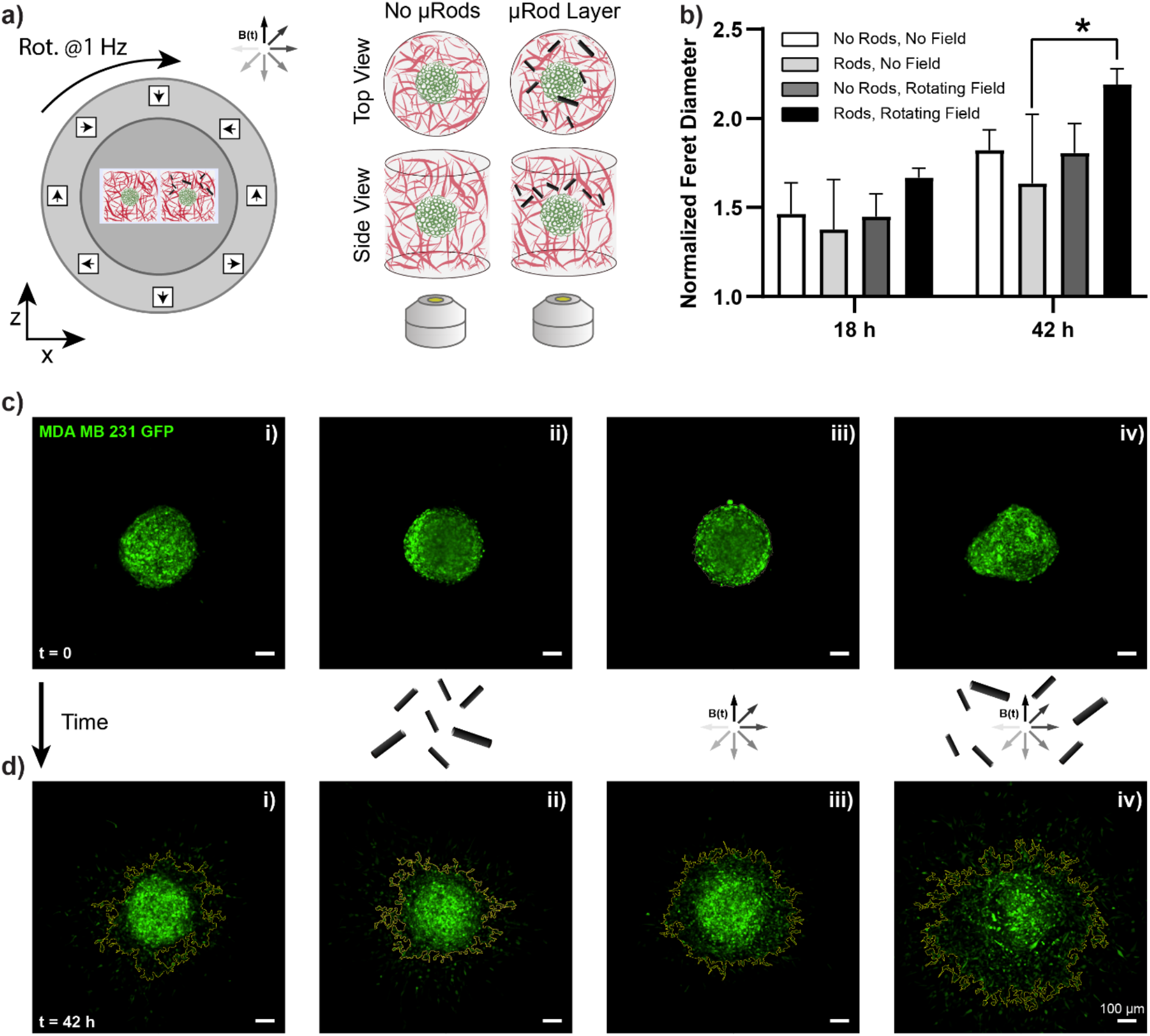
Tumor cell invasion is increased by cyclic microdeformation of the surrounding ECM. a) MDA-MB-231 GFP tumor spheroids embedded in Col I hydrogels at a concentration of 2 mg/mL were supplemented with a layer of Col I enriched with iron μRods. Samples were exposed to a rotating field of 50 mT magnitude under standard cell culture conditions and imaged after 18 h and 42 h of actuation. b) Invasion of cancer spheroids is quantified using the Feret diameter obtained from the analysis of fluorescence images acquired from the tumor spheroids and normalized with values obtained from images acquired 1 h after spheroid embedding in Col I. The graph shows mean and standard deviation. After 42 h of actuation, a significant increase in Feret diameter could be detected for spheroids cultured in the presence of actuated μRods compared to samples that were supplemented with μRods but not exposed to rotating magnetic fields. A Tukey’s multiple comparisons test and ANOVA was performed to test significance. If not indicated otherwise: ns; * indicates p<0.0118. (n=3). c)-d) Representative examples for the tested conditions (i) no field, (ii) μRods added, (iii) exposure to rotating magnetic field (50 mT, 1 Hz) without μRods, (iv) exposure to rotating magnetic field with μRods present. c) Spheroids embedded in Col I hydrogels for 1 h, before start of actuation experiments. d) The same spheroids as shown in c) were imaged after 42 h of incubation with or without exposure to rotating magnetic fields. The yellow outline indicates the outline of spheroid and was extracted using FIJI. All scale bars: 100 μm.

### Local Application of Cyclic Strain Triggers Anisotropic Invasion

To test the effect of localized micromechanical deformation, an experiment was executed analogously to the assay presented in the previous section with spatially confined µRods embedded at one side of a tumor spheroid. Thus, instead of the global distribution of μRod-enriched Col I hydrogel volumes across the whole surface of the spheroid sample, the suspension was applied only locally in a region adjacent to a spheroid (**Figure 6a**). In this manner, mechanical actuation of the Col I matrix could be spatially focused and only cells in close proximity to the applied μRods were directly exposed to the micromechanical deformation over time. We hypothesized that the cells would respond in a spatially dependent manner to mechanical stimulation and acquired images before actuation at t = 0 h, and 18 h and 42 h later. To quantify anisotropic response, images of spheroids were divided into two regions: one half associated with the hemisphere with locally applied µRods and an opposite hemisphere without µRods, and thus, no exposure to mechanical deformation (see also Supplementary Figure 9). Next, we anaylzed and compared the cell count within stimulated and unstimulated hemispheres (**Figure 6b**). Less pronounced increase in cell invasion between the hemispheres was observed for spheroids that did not experience actuation, in contrast to spheroids exposed to RMF that showed a strong anisotropic increase in cell invasion after 42 h. Representative fluorescence and bright field images of tumor spheroids with locally applied µRods with and without actuation are shown in **Figure 6c and d**.

**Figure 6:**
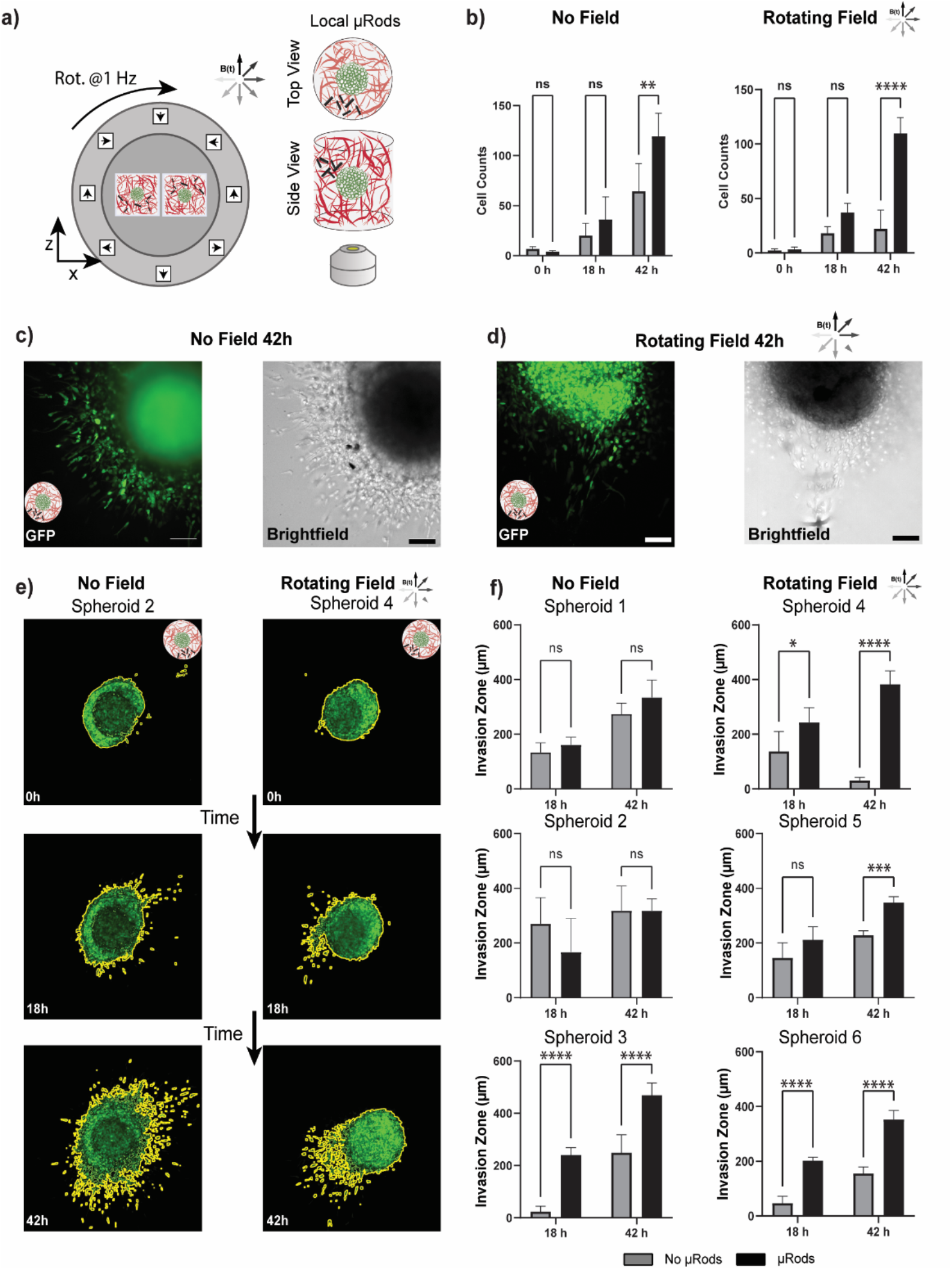
Investigation of tumor cell invasion in response to local actuation. a) MDA-MB-231 GFP tumor spheroids embedded in Col I hydrogels at a concentration of 2 mg/mL were supplemented with a layer of Col I and functionalized iron μRods were applied as local actuators on the periphery of the tumor spheroid. b) Cell invasion was quantified through dividing the images into two zones – with (black) and without (grey) µRods. A significant increase in cell counts was recorded in actuated spheroids in regions with µRods compared to regions of the opposite hemisphere (Two-way ANOVA, p < 0.0004). A less pronounced increase in cells was observed in spheroids not exposed to actuation. c and d) GFP and brightfield examples of rod positioning to spheroids with no field exposure and field exposure at 42h after embedding into Col I hydrogels respectively. e) Spheroids embedded in Col I hydrogels were imaged at time points 0, 18 and 42 h, after exposure to RMF of 50 mT magnitude under standard cell culture. All scale bars: 100 μm. f) The invasion zone of each spheroid was calculated within regions with and without µRods. The radius of the primary tumor was subtracted from distance of the furthest five cells from the primary tumor. Spheroid 3 was the only non-actuated spheroid that had a significant increase in invasion zone between rod and no-rod regions. All actuated spheroids had an increased in invasion zone between rod and no-rod regions at 42 h. A Turkey’s multiple comparisons test and ANOVA was performed to test significance. If not indicated otherwise: ns indicates p>0.1234 (not significant); * indicates p<0.0332; ** indicates p<0.0021; *** indicates p<0.0002; **** indicates p<0.0001 (n=3).

Overall, a trend of cell migration toward the location of the actuated μRods was observed. A particularly distinct example is shown Figure 6e (left), where the spheroid was exposed to RMF and cell migration occurs most intensely in the 7 o’clock direction, which is in the direction of the μRods (all analyzed spheroids are shown in **Supplementary Figure S9**). The extent of the invasion zone of the tumor spheroid into the Col I matrix was more closely investigated by measuring the distance of the five leading invading cells from the tumor rim (**Figure 6f, Supplementary Figure S9**). All spheroids exposed to RMF showed a significant increase in invasion zone distances in regions with µRods compared to no-µRod regions at latest after 42 h. We did, however, also observe one spheroid which had no exposure to RMF with apparent increase in invasion zone distance for the region with µRods versus without. This could have been triggered through mechanical perturbations during the addition of the μRods.

### Discussion and Conclusion

Cells and their surrounding microenvironment exist in a dynamic relationship, regarding the exchange of both biochemical and biomechanical signals. In particular, the process of cancer cell invasion is marked by the close interaction between cells and the surrounding ECM network. This work presents the controlled application of long-term mechanical deformation within Col I hydrogels that were used as model systems for studies of cancer cell invasion. Controlled by RMF, iron μRods served as embedded microscale actuators to achieve local deformation of the surrounding fibrous matrix.

Using particle imaging velocimetry analysis, strain rates within the hydrogel networks were analyzed with regards to magnitude and range of influence within the examined sample environments, showing regions of influence spanning 200 µm or larger, which is consistent with cell-generated strain fields. In long-term actuation experiments over 42 h, we showed that invasion of MDA-MB-231 cells from spheroids embedded in Col I hydrogels increased upon cyclic deformation of the Col I matrix by globally distributed μRods. Local application and actuation of μRods on one side of the tumor spheroids showed a local influence on cell migration, indicating local attraction of the invasive cancer cells by the localized strain.

By means of magnetic micro-actuators that were stably attached to Col I hydrogel matrices and showed good compatibility with cell culture systems (**Supplementary Figure S4**), strain was applied within a 3D Col I network for 42 h. The analysis of Col I displacement displayed in Error! Reference source not found. and Error! Reference source not found. revealed the rates of strain expressed in the Col I network during one cycle of actuation. The patterns of positive and negative values reflect on the direction of the strain and show alternation during one cycle of actuation, as to be expected. This indicates that during one cycle of actuation, the surrounding of one μRod experiences different directions of Col I fiber displacement and applied strain, resulting in a mechanically highly dynamic environment. The strain rates recorded through particle velocimetry analysis were dependent on the mode of actuation (in-plane or out-of-plane). This anisotropy is likely accounted by the more pronounced volumetric contribution of under lying planes for out-of-plane actuation. Certainly, the local structure of the Col I hydrogel might have also favored the higher rate of strain achieved by different degrees of μRods deflection within the surrounding network in this particular case and the strain fields will vary naturally for every µRod, as it does for therein migrating cells. Furthermore, the displacement of fluorescent tracer particles during μRods deflection out-of-plane might have influenced the signal recorded by fluorescence microscopy with regards to the focal plane. In this manner, higher degrees of displacement might be optically detected for the out-of-plane rotation of the magnetic field.

Col I is a biomaterial that is characterized by strain stiffening. The exposure of mechanically sensitive cells to these cyclically strained materials during magnetic actuation is therefore particularly interesting with regards to the temporal aspect of mechanical alteration of the cell surrounding matrix. Indeed, Ca^2+^ imaging revealed increased Ca^2+^ levels within cells exposed to sustained actuation of the surrounding matrix and increased invasion was observed for tumor spheroids of the invasive cell line MDA-MB-231. This partially supports previous findings by Menon *et al*. in 2011, reporting on increased invasion of the invasive sarcoma cell line HT1080 in response to cyclic strain resulting from magnetically controlled displacement of magnetic beads.^21^ However, Menon *et al*. attributed this increase of invasive behavior to a mechanism dependent on the presence of fibronectin as additional ECM component for the reception of the involved mechanical signals. In contrast to these findings, the presented system of Col I hydrogels does not require the addition of fibronectin to mechanically provoke increased invasive behavior. Potential reasons might include the investigated cell line, different patterns and magnitudes of magnetic particle actuation or different quality of the applied Col I matrix. Alternatively, the mechanisms underlying the change of invasion might differ with regards to the magnitude of the strain caused by deflecting μRods and the “tugging” forces described by Manon *et al*. While the displacement of beads resulting in the described “tugging” forces that enhanced cancer cell invasion in an integrin-dependent manner, the strains caused during the deflection of μRods might evoke strain stiffening of the Col I network, targeting stiffness-dependent modes of migration.^41^ Potential approaches to analyze exact mechanisms regulating the modes of migration could be investigated by RNA sequencing which offers the opportunity for unbiased screening of alterations in cellular regulation.

The potential effects of constant deformation over longer times might include a change in patterns of Col I architecture resulting from altered interaction of residing cells with their surrounding matrix. Further, the ECM is known to exhibit local accumulation of metabolites and signaling factor reservoirs that can be released through mechanical deformation by cells. A potential consequence of long-term cyclic deformation of the ECM network might influence the accumulation of such reservoirs and for instance result in the release of otherwise bound molecules and thereby causing changes in the biochemical environment of the tissue model.^42^

The presented work applies magnetic μRods that are stably attached to a Col I hydrogel matrix as sample-embedded microscale actuators. The design of a magnetic field generator that is suitable for use in a standard cell culture incubator allows for the actuation of several sample replicates under uniform conditions over several days of culture. Applied in the context of a 3D model for cancer invasion, the combination of techniques applied in this study is shown to influence the invasive behavior of cancer cells and to direct cell migration. While demonstrated on one cancer cell line, future studies could employ the presented techniques for the evaluation of mechanobiological influences in the context of other model systems and perform further readouts of biological markers.

## Author Contributions

Conceptualization: DA, SS; Data curation: DA, AM, NH; Formal Analysis: DA, AM, NH, RW, FL, SS; Funding acquisition SS; Investigation: SS; Methodology: DA, MC, SS; Project administration: SS; Resources: NA, SS; Software: DA, AM, AS, ADM; Supervision: SS; Validation: DA; Visualization: DA, RW, FL; Writing – original draft: DA, SS, AS, RW; Writing – review & editing: all authors;

## Acknowledgements

The authors would like to thank Prof. Salvador Pané, Siyu Deng, Fabian Landers and Valentin Gantenbein for technical support regarding the μRod fabrication and access to their electroplating setup. They also thank Cameron Forbrigger for proof reading and the staff of the FIRST clean room and of the Physics workshop at ETH Zurich for support.

## Supplementary Information

### Supplementary Text 1: Fabrication of Magnetic Iron μRods

Magnetic iron μRods were fabricated using template restricted electrodeposition. The method to prepare plating templates via photolithography was adapted from work by B. Özkale.^1^

### Preparation of Plating Templates

For the preparation of plating templates, 4-inch Si-wafers were coated with a conductive layer of 25 nm titanium and 125 nm gold using electron-beam evaporation (MEB550S, Plassys). The wafer was pre-baked at 200 °C for 10 minutes to remove any hydration layer on the surface and spin coated with a first layer of photoresist (10XT, 520 cp, MicroChemicals) at 500 rpm for 10 seconds followed by 2’400 rpm for 60 seconds. The wafer was soft baked at 110 °C for 80 seconds and left to rehydrate for 10 minutes. Next, a second layer of photoresist was applied using previously listed spin parameters and soft baked at 110°C for 160 seconds. For rehydration, the coated Si-wafer was stored at ambient conditions for 45 minutes prior to exposure. Exposure of the photoresist was performed using a mask aligner (EVG 620 NT, EVG) with a chrome mask designed for arrays of pores with a diameter of 5 μm in a distance of 15 μm (Jena masks, Germany) at a dose of 400 kJ/cm^2^.

Post exposure, samples were developed using AZ 400 K developer (MicroChemicals) at a dilution of 1:3 for 17 minutes under gentle agitation and exchange of developer after 10 minutes of incubation. Samples were rinsed in excess with deionized water and dried with pressured air. Si-wafers were diced into chips of 2*3 cm^2^ for plating experiments using a diamond scribe.

### Plating of μRods

Plating of iron μRods was performed using photoresist-based templates which were prepared as specified in the previous section. The plating templates were subjected to oxygen-plasma treatment for 30 seconds prior to electroplating experiments to enhance surface wetting.

The electrodeposition methodology was adapted from Alcantara et al.^2^ The electrolyte solution was prepared with amounts as specified in **Supplementary Table 1**. Deionized water was added to reach a final volume of 200 mL and the solution was stirred at 250 rpm with a magnetic stirrer at 50°C to obtain a green transparent electrolyte solution. Concentrated sulfuric acid (Sigma-Aldrich) was added dropwise to adjust the solution to pH 2 and the stirring frequency was reduced to 50 rpm for the plating experiments. Plating templates were connected to the working electrode and immersed in the electrolyte solution together with a 5*5 cm platinum sheet counter electrode and Ag/AgCl reference electrode. A potential of −0.95 V against the reference electrode was applied for 40 minutes to obtain iron μRods with an approximate length of 25 μm. After plating, samples were removed from the electrolyte solution, rinsed with deionized water, and dried with pressured air.

The photoresist template was removed by thorough washing with acetone and isopropanol, gently dried with pressured air. A protective layer of silicon oxide (SiOx) with a thickness of approximately 60 nm was applied via plasma enhanced chemical vapor deposition (PECVD 80+, Oxford Instruments). The SiOx-layer provided a bioinert shell around the μRods to prevent corrosion and the leakage of iron or iron oxides into the surrounding aqueous environment.

### SEM Imaging of μRods in Col I

For SEM imaging of Col I hydrogels enriched with iron μRods, Col I hydrogels were prepared as specified above and supplemented with 5 μL of surface-functionalized iron μRods. Hydrogels were pipetted on round pieces of filter paper and left to polymerize at 37°C, 5% CO2 in a standard cell culture incubator for 45 minutes. Samples were fixed for 1 hour in glutaraldehyde, thoroughly washed with deionized water and dehydrated by means of serial incubation with increasing concentrations of ethanol. Samples were subjected to critical point drying and sputtered with Pd/Au prior to imaging.

### Surface Functionalization of Iron μRods

A surface functionalization of the SiOx-coated iron μRods was performed to allow for stable attachment of the iron μRods to the surface of Col I hydrogel networks via amidation. To stably link the surface of SiOx-coated iron μRods to the fibrous network of Col I hydrogels, surface of the coated μRods was silanized to obtain free amine groups (NH2) on the particle surface. Next, an NHS-PEG-NHS linker was grafted onto the NH2 groups. The functionalization strategy is depicted in **Supplementary Figure S1** and is described in the following paragraphs.

### Amine Functionalization of SiOx-Coated μRods

The silanization process of the SiOx-coated iron μRods was adapted from Kim et al.^3^ Clean and dry SiOx-coated sample chips were placed in 50 mL Falcon tubes and covered with a solution of 2% 3-Aminopropyltriethoxysilane (APTES, 99%, Acros) that had been prepared with anhydrous toluene (extra dry, Fisher Scientific). Samples were incubated for 4 hours at room temperature under gentle agitation, rinsed two times for 10 minutes with toluene and dried with nitrogen. Until further use, samples were stored in filtered deionized water at 4°C.

### Grafting of NHS-PEG-NHS on the Surface of Amine-Functionalized μRods

For stable attachment of iron μRods to free amines of Col I networks, NHS-groups were grafted on amine groups of silanized μRods according to a previously published method.^4^ Amine-functionalized μRod chips were incubated with a solution 5 mg/mL NHS-PEG-NHS (4-arm PEG, 5k, Creative PEG Works) in TEA buffer (0.1 M, pH 6) and incubated at RT under gentle agitation for 30 minutes. Samples were washed twice in TEA buffer (0.1 M, pH 6) and sonicated for a few seconds to release functionalized μRods from the plating substrate.

### Experimental Verification of μRod Surface Functionalization

Silanized and non-silanized μRods were incubated with 5(6)-TAMRA *N*-succinimidyl ester (C0027, Chemodex, hereafter referred to as NHS-TAMRA) diluted at a concentration of 5 mg/mL in 0.1 M TEA buffer (pH 6) for 30 minutes under gentle agitation at room temperature and thoroughly washed twice with 0.1 M TEA buffer (pH 6). Samples were briefly sonicated to disperse the μRods.

Drops of μRod suspension were pipetted on a microscopy slide and imaged via fluorescence confocal microscopy (Nikon Eclipse) in the TEXAS Red channel (580 nm) (**Supplementary Figure S2**), confirming the attachment of the NHS-TAMRA dye to the μRod surface via amidation. The control sample of non-silanized, SiOx-coated μRods, which had been incubated with the same dye, did not exhibit a signal, confirming that the fluorescent labeling of the silanized sample was not caused by unspecific binding of the dye to the μRod. Similarly, the successful grafting of NHS-groups onto the silanized μRod surface was experimentally confirmed.

Following the functionalization of μRods with NHS-PEG-NHS groups as described before, μRods were incubated with an Amine-Cy3 dye (Cyanine3-amine, Lumiprobe) that was diluted in 0.1 M TEA buffer (pH 6) at a concentration of 5 mg/mL for 30 minutes under gentle agitation at room temperature. As a control, NHS-PEG-NHS functionalized μRods were simultaneously incubated with a 50 M Tris-acetate-EDTA (TAE) buffer to quench any reactive NHS sites with an excess of amines present in the buffer. Next, both the quenched control and the Amine-Cy3 labelled μRods were washed with 0.1 M TEA buffer (pH 6). Then, to test for unspecific binding of the dye, the Amine-Cy3-labelled μRods were incubated with the previously described quenching solution while the NHS-quenched μRods were incubated with the previously specified Amine-Cy3 labeling solution. In both cases incubation lasted 30 minutes. Subsequently, samples were rinsed twice with 0.1 M TEA buffer (pH 6) and sonicated to disperse the μRods. Drops of the μRod suspensions were pipetted onto microscopy slides and imaged via confocal fluorescence microscopy to test for fluorescence signal. A strong signal of Amine-Cy3 labelling was detected for the NHS-PEG-NHS-functionalized μRods while the quenched samples do not exhibit a clear fluorescence signal (**Supplementary Figure S2c**), indicating that the quenching procedure blocked all available NHS-groups prior to the fluorescent labeling and that the unquenched samples exhibited reactive NHS-groups that were well reacted with the amines of the Amine-Cy3 dye.

### Magnetic Characterization of Iron μRods

SiOx-coated μRods were magnetically characterized via vibrating sample magnetometry (VSM). To prepare the sample, Polydimethylsiloxane (PDMS) polymer (SYLGARD^TM^ 184 Silicone Elastomer, Dow) was mixed according to manufacturer’s instructions and degassed for 1 hour. 1.5 mg of SiOx-coated μRods were dispersed in 100 μL degassed PDMS that was then poured in a mold ring of cured PDMS with 8 mm diameter that had been previously prepared in a plastic dish. The sample was degassed again for 1 hour at room temperature and then cured at 80°C for 3 hours. Using a metal puncher with 8 mm diameter, the PDMS containing magnetic μRods was punched out of the template. A background control sample was prepared accordingly without addition of magnetic μRods.

For VSM measurements (**Supplementary Figure S3**), sample disks were glued to the sample holder (8 mm Pyrex Transverse) using double sided tape. A hysteresis loop was measured using a vibrating sample magnetometer (EZ-VSM, Microsense). The background signal from the sample holder and blank PDMS sample was measured and subtracted from the signal collected for the μRod-containing sample.

### Biocompatibility Analysis of Iron μRods

Biocompatibility of functionalized iron μRods was tested via an MTT assay (**Supplementary Figure S4**). Cells of the invasive cancer cell line MDA-MB-231 GFP cells were cultured as described above and were seeded in 96-well plates at a density of 10^4^ cells/well. After one day of culture, 97 μL of the respective cell culture medium were added together with functionalized μRods dispersed in 3 μL of 0.1 M TEA in two different amounts (Low Concentration: 50 μRods per well; High Concentration: 250 μRods per well). Further, for the vehicle control, 97 μL of culture medium were supplemented with 3 μL of 0.1 TEA buffer without μRods and an untreated control was provided with 100 μL of fresh cell culture medium. All samples were prepared in triplicate and analyzed via the MTT assay 1, 2, and 3 days after μRod addition. Cell viability was quantified using the CyQuant MTT Cell Viability Assay (Thermo Fisher Scientific, V13154). After 10 minutes of cell incubation with the reaction mix at 37°C and 5% CO2, samples were gently agitated and a multimode microplate reader (Spark, Tecan) was used to measure absorbance at 540 nm. The experiment was performed using triplicates.

### Supplementary Text 2: Image Processing

Imaging of spheroids embedded in Col I hydrogels was performed with 10x magnification and image stacks were acquired with 8 μm distance and a total of 11 images per stack. During imaging, care was taken to center the equatorial region of the spheroid in the center of the stack, to capture the maximum spread and flanking regions. Imaging of spheroids was performed 1 hour after embedding, and after 18 hours and 42 hours of incubation.

If not indicated differently, acquired images were processed using FIJI. Analysis of spheroid morphology in Col I hydrogels was performed the following way using FIJI: Z-projections of image stacks acquired of green-fluorescent tumor spheroids were generated and saved as .tiff files. All images were subjected to the same adjustment regarding window level, converted to binary files, thresholded and analyzed using the particle analyzer function of FIJI. Parameters extracted from the image data included projected area and centroid point of the primary tumor and its subsequent invading cells, the minimal and maximal Feret diameter, including the x and y axis of the spheroid, depending on the intended analysis. The invasion zone of each spheroid was calculated by measuring the distance (μm) of the top five furthest travelled cells from the rim of the tumor.

### Supplementary Text 3: Explanation of Torque Estimation for µRods

The soft ferromagnetic µRods employed in this study consist of metallic iron and are large enough to support multiple magnetic domains. Under the application of an external magnetic field such as the 50 mT field used in actuation experiments, they can be regarded as approximately uniformly magnetized and experience anisotropy dominated by shape effects, with their uniaxial easy axis coinciding with the axis of the cylinder. At low fields, the magnitude of the torque will scale proportionally with the moment of the µRod 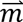 and the applied field 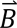 with its upper limit 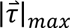 given by

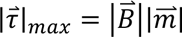

For the purpose of estimating this value for the µRods used in this work, we can assume an applied field magnitude of 50 mT, a µRod length *l* of 25 µm, a µRod diameter *d* of 7 µm, and a saturation magnetization *M_S_* corresponding to that of bulk metallic iron.^5^

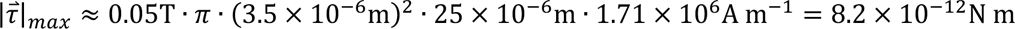

During cyclic actuation by rotating magnetic fields, this maximum torque would only be reached briefly during the part of the cycle when the moment is perpendicular to the field. It should additionally be noted that the absolute upper bound of available torques that can be supplied with a µRod of this type is set by the energy scale of the magnetic shape anisotropy. In the limit of high applied fields, the moment would be oriented away from its easy axis, but a torque resulting from shape anisotropy would still be experienced. The estimate employed above assumes that the moment is still confined to the easy axis under the applied field, an assumption that can be supported by comparing that torque value to that of the ultimate torque available from shape anisotropy. The magnetostatic energy *U* of a magnetized cylinder with its moment oriented at angle θ relative to the axis of the cylinder is

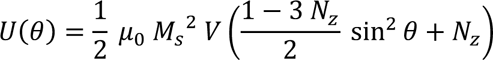

The demagnetizing factors for cylinders, *N_x_* = *N_y_* and *N_z_*, expressed as a function of their aspect ratio *n*, where *n* = *l*/*d* (length over diameter), have been computed numerically and can also be given by approximate analytical functions.^6^ The torque τ resulting from this potential energy function is

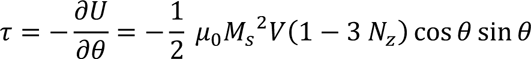

With the dimensions assumed above, the aspect ratio is approximately 3.6, giving an *N_z_* value of approximately 0.11. Thus, for the µRods employed in this work, the absolute maximum applicable torque is estimated to be

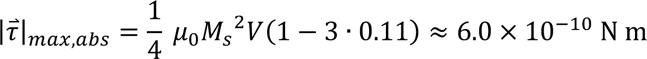

Because this value is approximately two orders of magnitude higher than the torque estimated using just the applied field and the moment of the µRod, it is clear that the earlier estimate assuming that the moment is largely confined to the easy axis is well motivated for the conditions we employed in this study.

**Supplementary Table 1:**
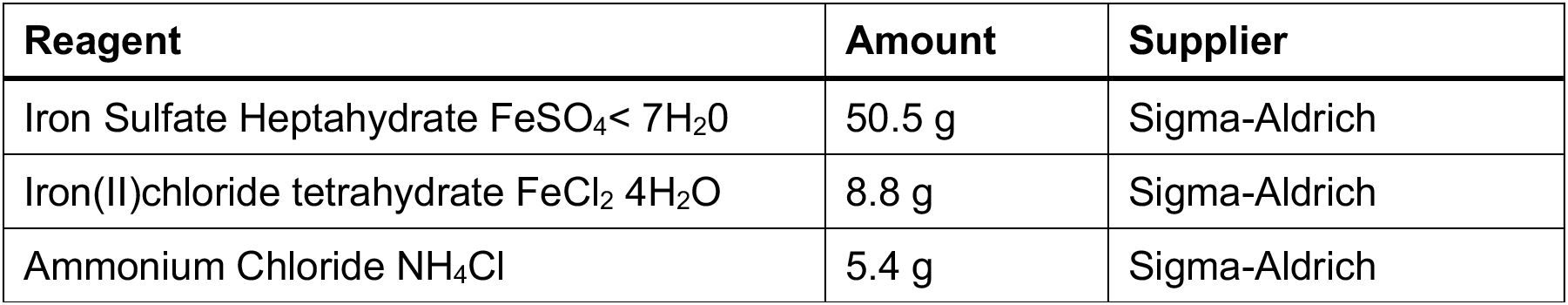
Composition of the electroplating solution The table lists the components for the preparation of an electrolyte solution for the electrodeposition of iron µRods

**Supplementary Figure S1:**
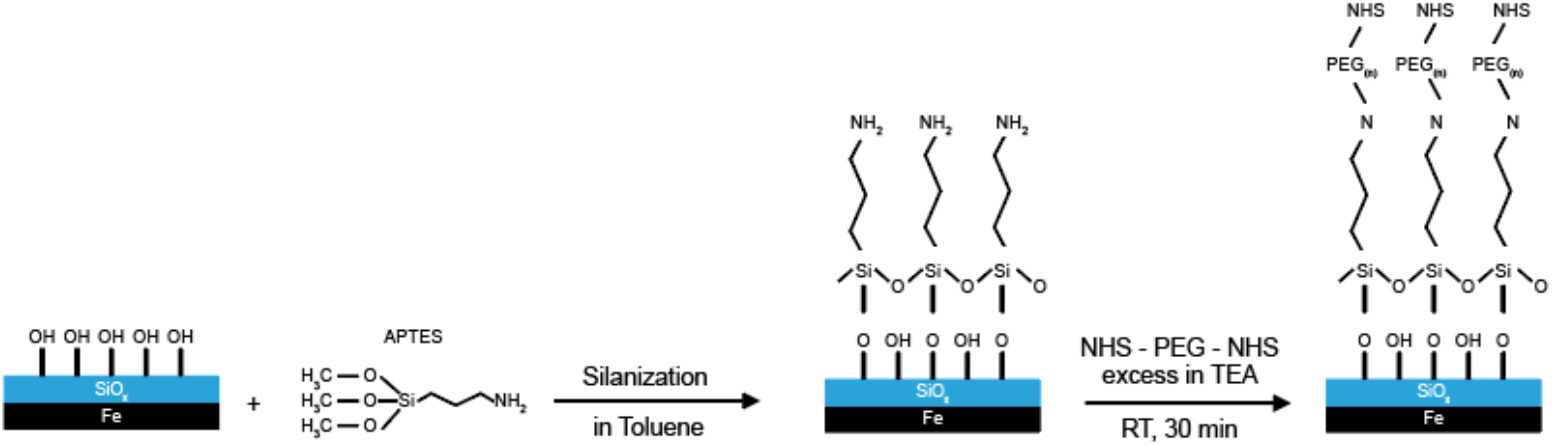
Amine functionalization of μRods. The schematic depicts different stages of the surface functionalization strategy of SiOx-coated iron μRods to obtain NHS-groups that can bind to free amines exposed by the Col I network via amidation. The illustrated process is described in the Materials & Methods section.

**Supplementary Figure S2:**
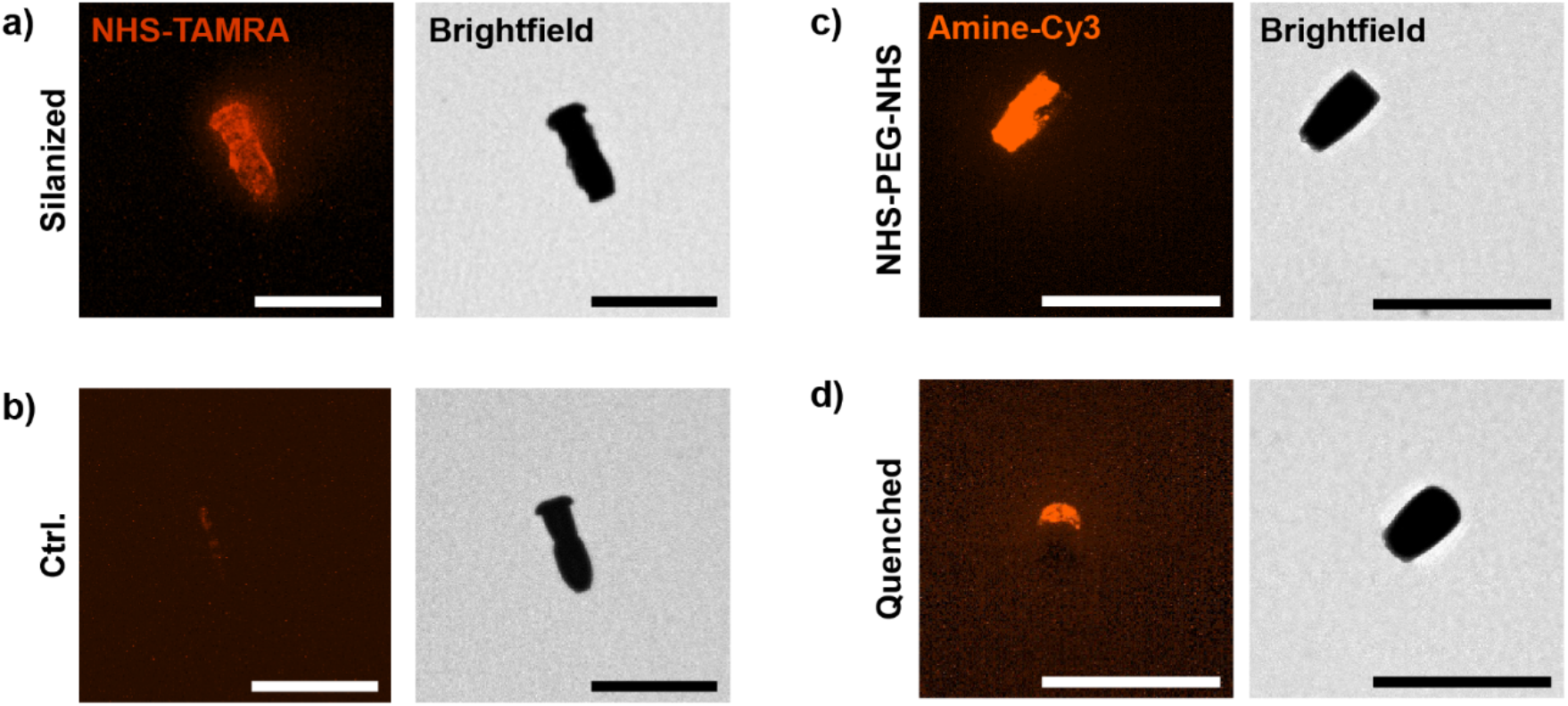
Experimental verification of SiOx μRod surface functionalization via fluorescence confocal microscopy. a-b) Silanized and untreated μRods were incubated with an NHS-TAMRA ester. **a)** Confocal fluorescence (left) and brightfield image (right) of a silanized μRod that was exposed to a solution of NHS-TAMRA to stain for free amines on the μRods surface. **b)** Confocal fluorescence (left) and brightfield image (right) of a non-silanized μRod that was exposed to the same staining solution as the sample shown in a). **c-d)** An Amine-Cy3 dye was used to test for the presence of reactive NHS-groups on the μRod surface after surface functionalization of the μRods with an NHS-PEG-NHS ester. **c)** Confocal fluorescence (left) and brightfield image (right) of a NHS-PEG-NHS-functionalized μRod labelled with an Amine-Cy3 dye, indicating the presence of active NHS groups. **d)** Confocal fluorescence (left) and brightfield image (right) of a NHS-PEG-NHS-functionalized μRod which had been quenched with a 50 M TAE buffer to block all present NHS groups prior to labeling with an Amine-Cy3 dye. All images are representative examples. All scale bars: 50 μm.

**Supplementary Figure S3.**
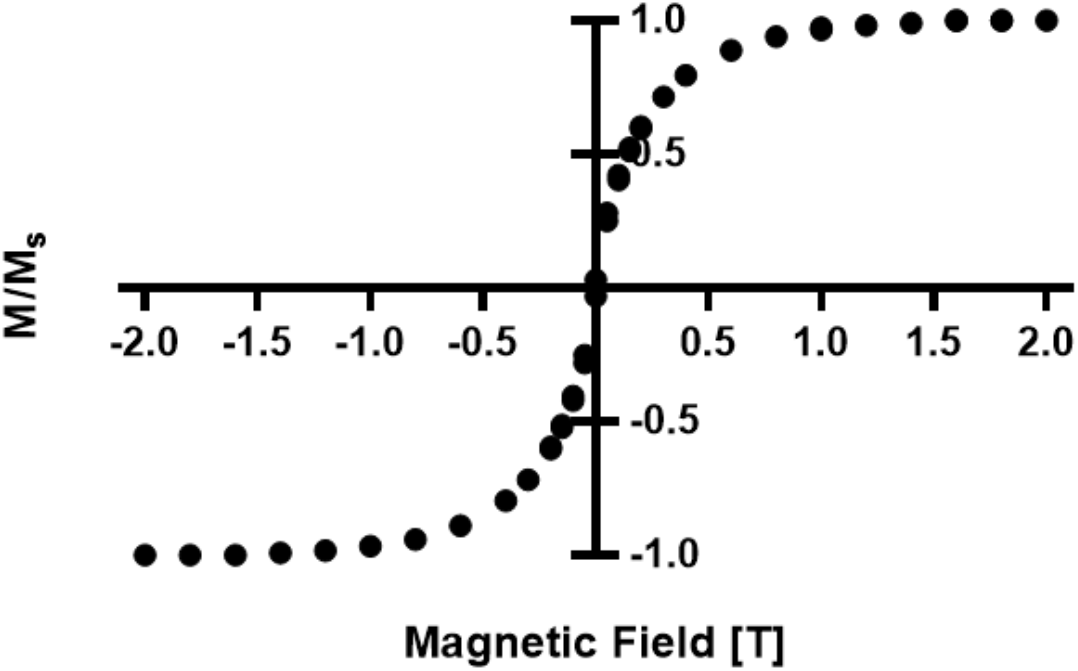
Vibrating sample magnetometry (VSM) analysis of iron μRods. The data from the VSM measurement confirm the ferromagnetic character of the iron μRods.

**Supplementary Figure S4:**
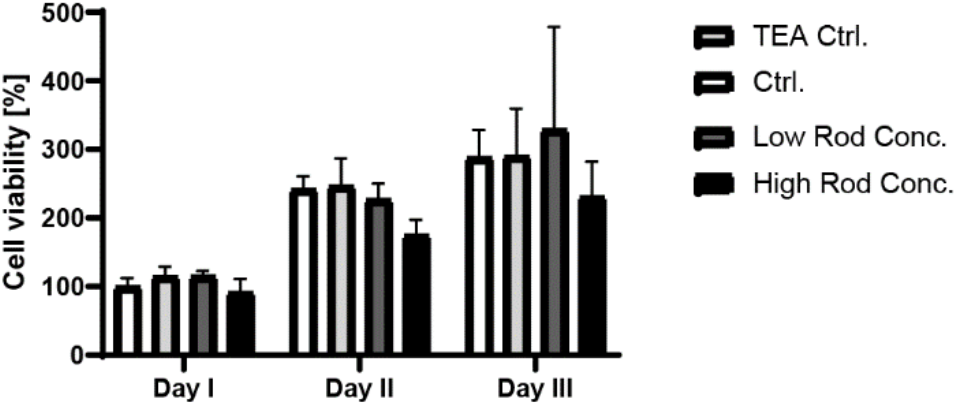
Biocompatibility analysis of surface functionalized iron μRods tested via MTT assay. Data is shown for MDA MD 231 GFP cells that were cultured in the presence of different μRod concentrations and tested for viability over three days of culture. Ctrl: No treatment; TEA Ctrl: 3 μL of 0.1 M TEA buffer without μRods; Low μRods Conc.: 50 μRods dispersed in 3 μL of TEA buffer per well; High Rod Conc.: 250 μRods dispersed in 3 μL of TEA buffer per well. (n=3) Significance was tested using the Turkey’s multiple comparison test and no significant difference was found between the conditions. The results suggest that no significant toxicity could be detected for neither the high nor low μRod concentrations.

**Supplementary Figure S5:**
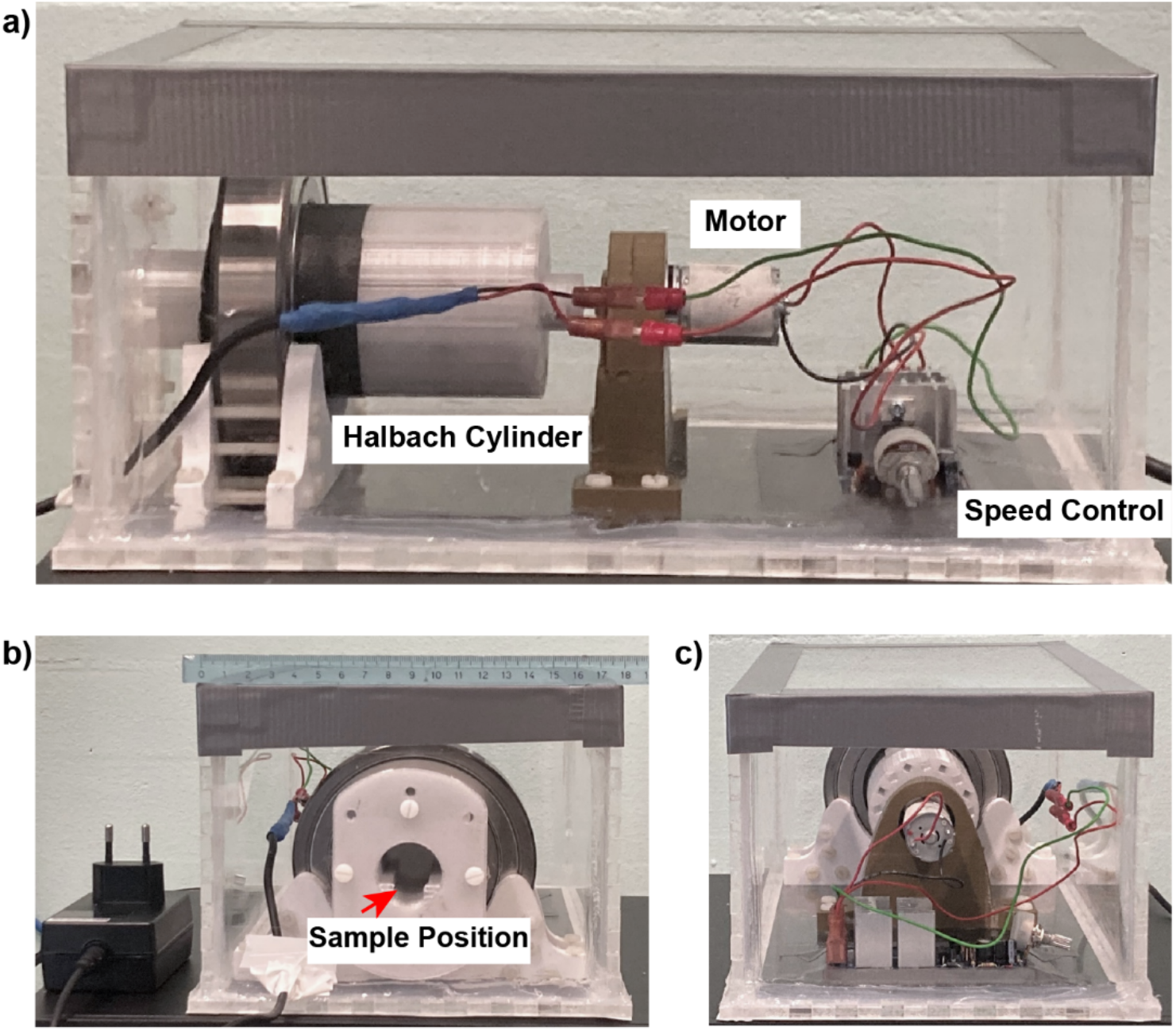
Overview of the Halbach-cylinder-based magnetic field generator. Photographs show the magnetic actuation setup which is encased in a sealed acrylic box. **a)** side view, indicating the position of the Halbach cylinder, the actuating motor and the speed controller. **b)** Front view of the setup, indicating the position where sample slides were introduced to the system for long-term actuation. **c)** Rear view of the setup.

**Supplementary Figure S6:**
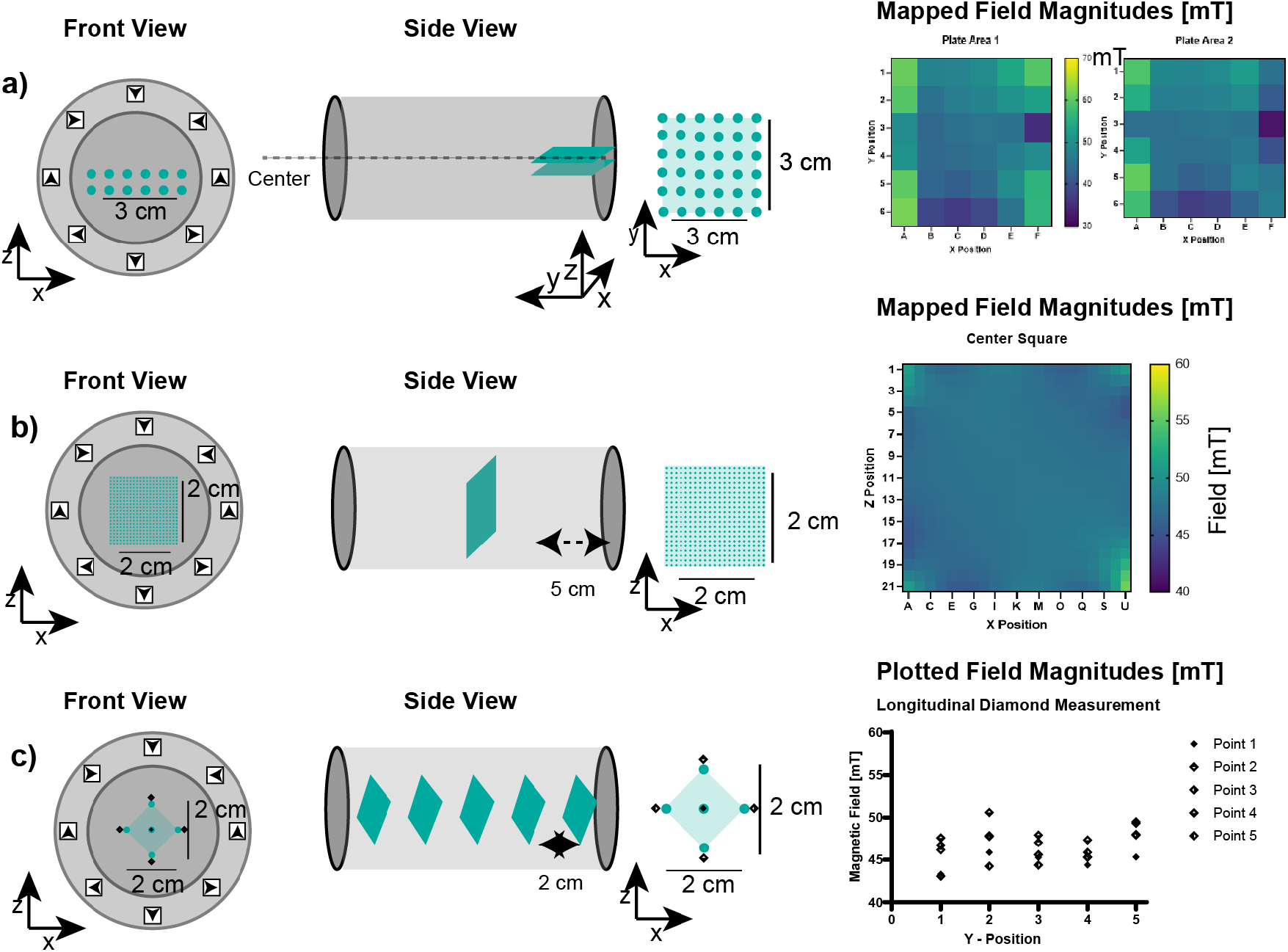
Magnetic field characterization of the Halbach cylinder. Different sets of measurements were collected to characterize the field magnitude within in the previously described Halbach cylinder. The left column of sketches displays the front view of the cylinder, in the center, the cylinder is depicted from the side. Blue dots indicate locations of individual measurement points. The right column of images displays color-coded maps of measured magnetic field magnitudes, displayed in mT. The analysis of the magnetic field characterization indicated a high homogeneity of the generated magnetic field. An approximate field magnitude 50 mT was achieved throughout the cylinder in center regions where the samples would be placed during experimental procedures. To incubate samples, small plastic containers containing PBS-soaked sterile tissues were placed next to the samples to avoid sample dehydration. Experiments without exposure to magnetic fields were incubated in humidified petri dishes. **a)** Two planes were mapped inside the cylinder at locations that marked the upper and lower edges of the applied cell culture ware. **b)** A square was mapped in the center of the cylinder to investigate homogeneity of the field magnitude. **c)** To investigate differences in magnetic field magnitude at positions in the center and towards the edge of the cylinder, repeated measurements were taken as indicated by the diamond-shaped array for different positions inside the Halbach cylinder.

**Supplementary Figure S7:**
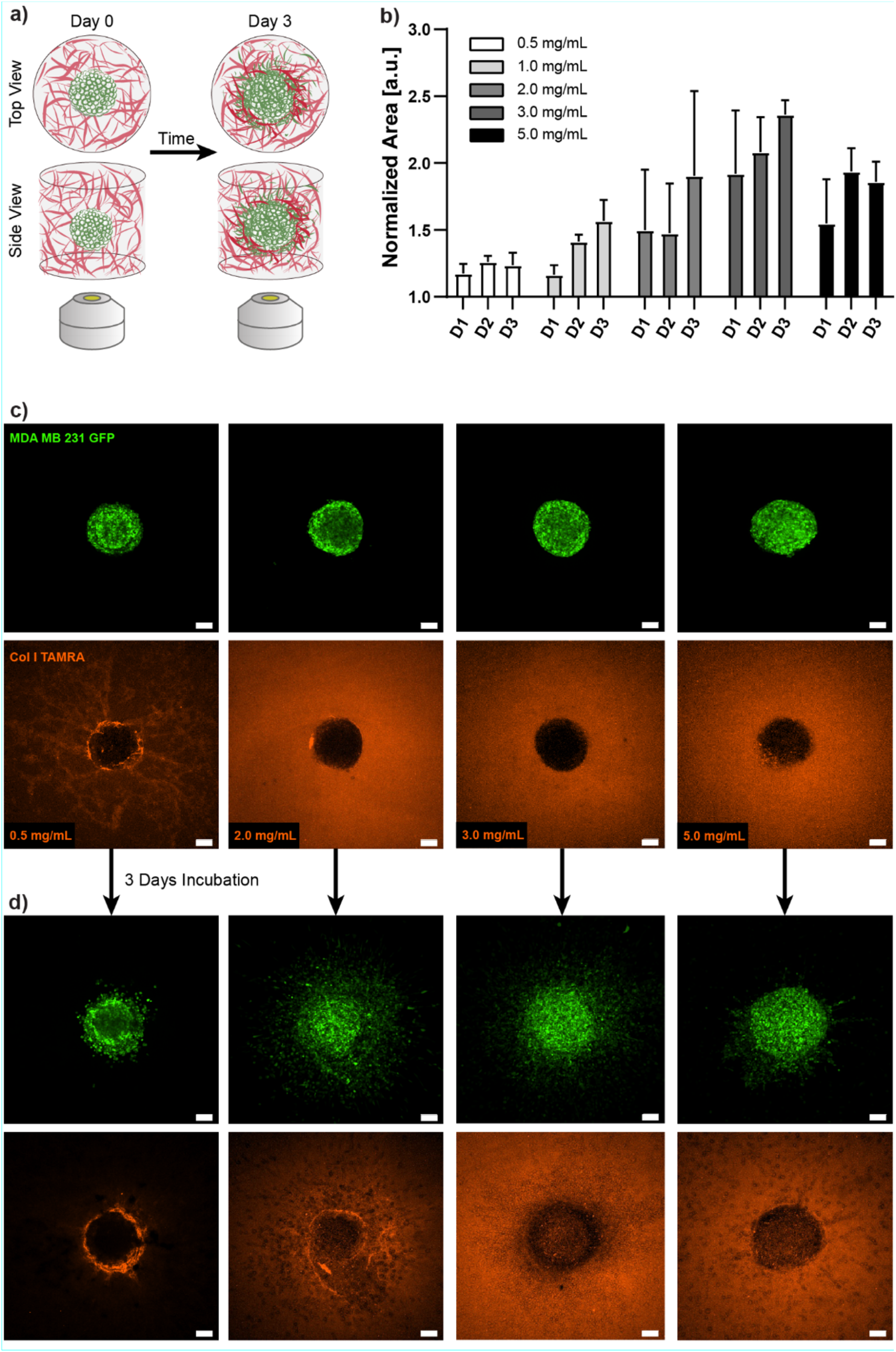
Concentration-dependent invasive behavior of invasive cancer cells to the surrounding extracellular matrix. **a)** Tumor cell invasion is modeled by embedding of tumor spheroids from the invasive breast cancer cell line MDA MB 231 (displayed in green) in Col I hydrogel matrices (labelled in red). Tumor cell invasion is investigated by repeated imaging and analysis of cellular spreading into the surrounding matrix network. (Indicated by the objective). **b)** Tumor cell invasion was tested for different concentrations of Col I hydrogels, between 0.5 and 5 mg/mL and analyzed for invasion of cancer cells via confocal fluorescence microscopy over three days of culture. The area of tumor cell spreading on fluorescence images was analyzed using FIJI and normalized with the area determined for images acquired 1 hour post embedding of the tumor spheroids. The graph shows mean and standard deviation. (n=4) **c)** Representative fluorescence confocal images as acquired and processed for the analysis of tumor cell invasion into the surrounding Col I hydrogel matrix. Examples for concentrations between 0.5 mg/mL and 5 mg/mL are shown. MDA MB 231 GFP cells are depicted in the top row, TAMRA-labelled Col I hydrogels are shown in red in the images of the bottom row. Data is shown for Day 0, imaged 1 hour post embedding of the tumor spheroids. **d)** Confocal fluorescence images of samples shown in c), imaged after 3 days of incubation. All scale bars: 100 μm.

**Supplementary Figure S8:**
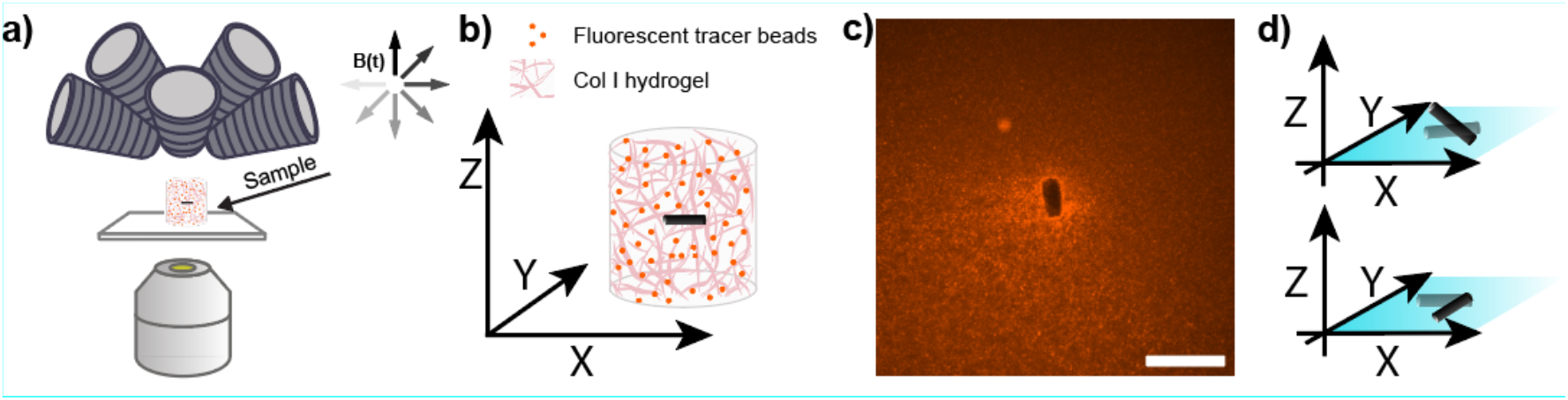
Experimental setup for the analysis of Col I hydrogel deformation in response to magnetic actuation of sample-embedded magnetic μRods. **a)** Samples were imaged during cyclic magnetic actuation using an eight-coil magnetic field generator. **b)** Fluorescent tracer beads (red) dispersed in the Col I hydrogels allowed to track Col I hydrogel deformation in response to magnetic actuation. **c)** Representative image of an iron μRod embedded in a Col I hydrogel (2 mg/mL) enriched with fluorescent microparticles (indicated in orange-red). Scale bar: 50 μm. d) Out of plane (top sketch) and in plane rotating magnetic fields (bottom sketch) were used to actuate Col I-embedded μRods to capture the 3D character of the sample volumes.

**Supplementary Figure S9:**
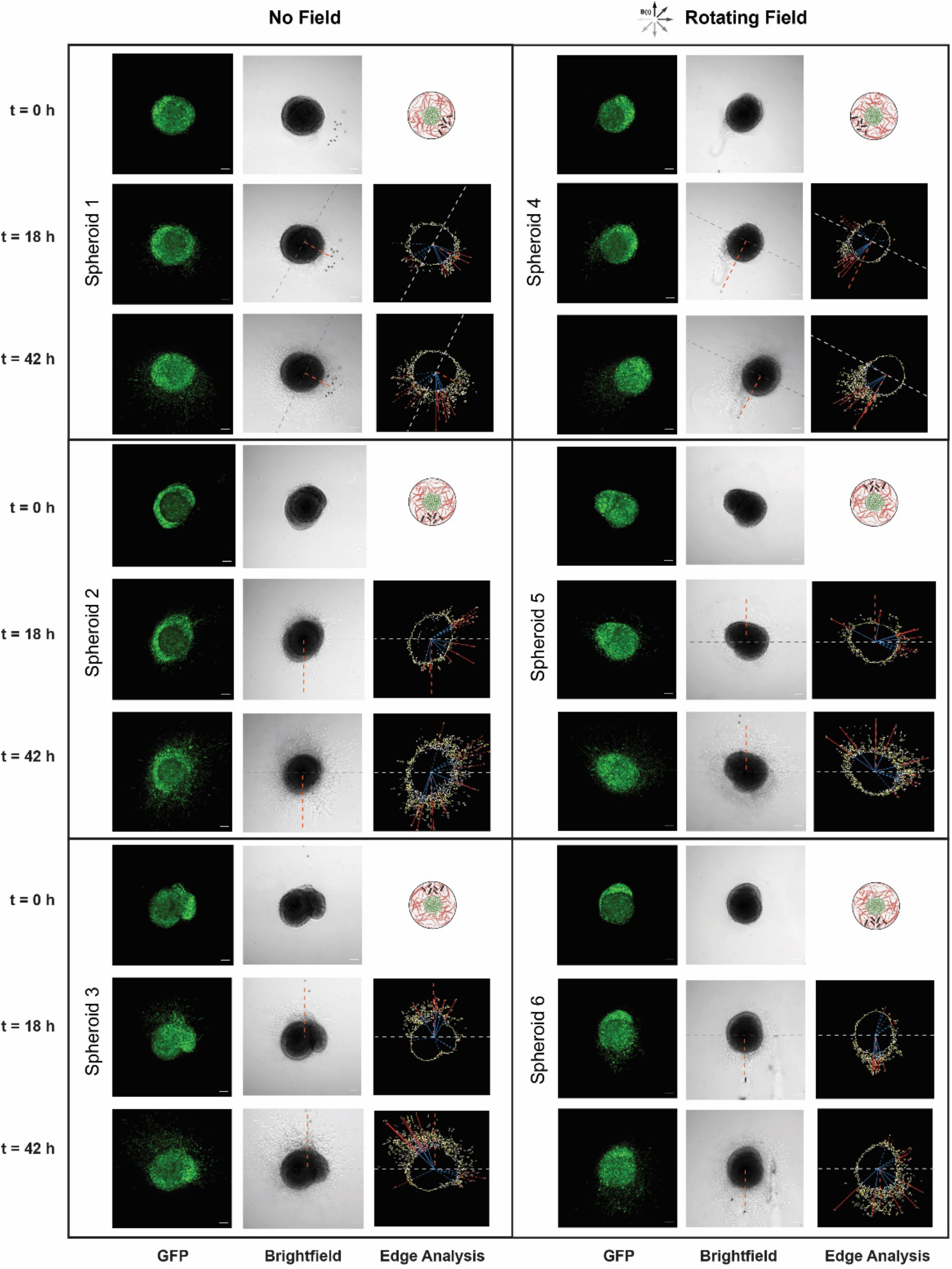
Investigation of tumor cell invasion in response to local actuation. MDA-MB-231 GFP tumor spheroids embedded in Col I hydrogels at a concentration of 2 mg/mL were supplemented with a layer of Col I and functionalized iron μRods were applied as local actuators on the periphery of the tumor spheroid, then imaged after 0, 18 and 42 hours, of incubation, respectively, and after exposure to RMF of 50 mT magnitude (or no field) under standard cell culture conditions. Edge analysis was performed for each spheroid to identify invasive cells (see Materials & Methods). Cell invasion was quantified through dividing the images into two zones (gray dashed line) – with and without µRods. The division line is derived by taking orthogonal line of relative position between μRods and the center of primary tumor (orange dashed line). In each zone, furthest five invasive cells were collected, and the invasion zone (red line) was calculated by subtracting the radius of the primary tumor (blue line) from distance of invasive cells from center of the primary tumor. All scale bars: 100 μm.

## Supplementary Videos

**Supplementary Video V1:** Microscopy recording with a 40x objective of embedded μRod actuation (50mT, 1Hz) in Col I networks during in-plane magnetic actuation to assess strain rates

**Supplementary Video V2:** Microscopy recording with a 40x objective of embedded μRod actuation (50mT, 1Hz) in Col I networks during out-of-plane magnetic actuation to assess strain rates

**Supplementary Video V3:** Ca2+ influx imaging during magnetic actuation

